## Supplementary material for "ZAF – The First Open Source Fully Automated Feeder for Aquatic Facilities": ZAF supplementary information (v2)

#### Supplementary informations

**Merlin Lange, AhmetCan Solak, Shruthi Vijaykumar, Hirofumi Kobayashi, Bin Yang & Loic A. Royer [1]**

[1] Chan Zuckerberg Biohub, San Francisco, USA

The latest version of this material can be found on the wiki https://github.com/royerlab/ZAF/wiki

- **Table of contents**


  - Introduction
  - ZAF - Hardware
    * Frame
    * Servo & food container
    * Pumps & Valve
    * Food and mixing container
    * Safety pump and sensor
    * Tubing
  - ZAF - Electronics
    * Wire connections
    * Motor drivers
    * Raspberry Pi and Servo Hat Board
    * Screen
  - ZAF - Software
    * Installation
    * How to use
  - Tips to scale up ZAF
  - ZAF equipments
  - ZAF+ - Hardware
    * Frame
    * Food container Servo
    * Pumps & Valve
    * Safety
    * Tubing
  - ZAF+ - Electronics
    * Safety and connectivity
    * Arduino Mega
    * Motor drivers
    * 16 Relay Module
    * Raspberry Pi & Screen
  - ZAF+ - Software
    * Installation
    * How to use
  - Tips to scale up ZAF+
  - ZAF+ equipments
  - Troubleshooting

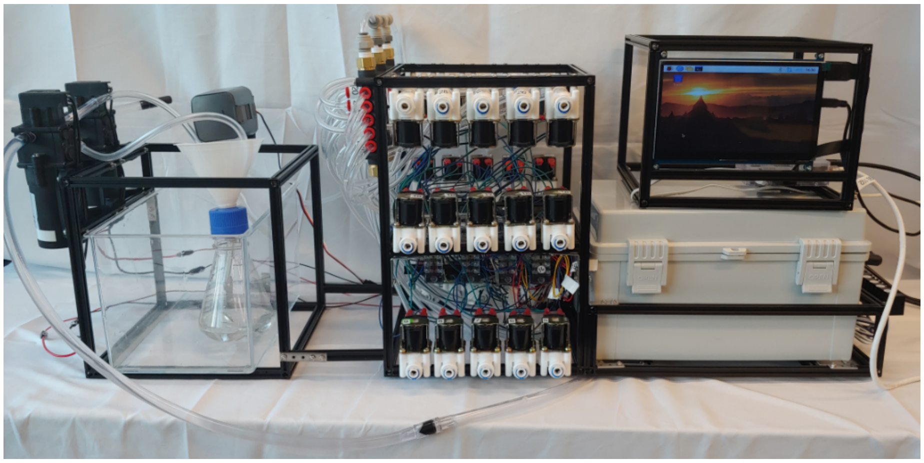

Figure 1: ZAF+

ZAF which stands for ‘Zebrafish Automatic Feeding’ is an easy to build device that can automate fish feeding for your fish facility with minimal supervision. ZAF is an open access project with both hardware and software designs openly provided. We give all instructions for building ZAF, leveraging only commercially available parts. ZAF is designed to be maximally accessible for non-experts in electronics, mechanics and robotics – we avoided complex parts, 3D printing, advanced electronics and tried to keep the design simple. We are providing a general manual to build ZAF, but the design is modular and can be adapted to the users need and usage.

ZAF comes in two versions: ZAF and ZAF+. The original ZAF is very easy to build but has some limitations (see below). ZAF+ addresses most shortcomings of ZAF but has a more advanced design but is also harder to build. The main limitation of ZAF is that it lacks precise control over the amount of food distributed to individual tanks. For example, ZAF distributes the same homogenised mix of food and water to all tanks. Therefore, if you have 12 fish in a tank and 6 in another, with ZAF they will receive the same amount of food. ZAF+ has the key advantage that it can control the amount of food you provide to each tank in your facility.

We have used ZAF successfully in our facility and found it to be very beneficial for general feeding during the weekend and holidays. However, if what you need is daily automated feeding then ZAF+ is a better choice. ZAF+ builds upon the existing modules of ZAF, but adds better flow control and variable food delivery thanks to more efficient pumps and a valve-based distribution system. Moreover, ZAF+ design allows for arbitrary scaling of the number of feeding lines, thus making it possible, in theory, to automatically feed fish in large facilities. We give instructions for both versions because we believe that different fish facilities and different users will have different needs and skills.

We designed a Graphical User Interface (GUI), to help the user run their own feeding programs. We tried our ZAFs with both dry and solid food and it works perfectly well. ZAF can help overcome staffing issues during weekends and holidays and can be used together with a camera system for remote monitoring. Additionally, ZAF+ can control the amount of food per tank depending on the fish number or density. In our manuscript, we show that ZAF and ZAF+ are good for fish health and reproduction rate and did not affect the water quality of the fish rack. All together, we hope that ZAF will help to facilitate the establishment of fish colonies in labs and will push zebrafish research into a new era.

You will find the detailed protocols to build your own ZAF and ZAF+. Both share a common design principle and ZAF+ is designed as an extension of ZAF. So, if after building ZAF you feel adventurous, we strongly recommend you build ZAF+, a slightly more complex device, but with more advanced features. Thus, even though we present two independent manuals, they share some common modules such as the food mixing.

The manuals for ZAF and ZAF+ are divided into three main parts:

- The hardware
- The electronics
- ZAF control software

The latest version of this material can be found on the wiki.

##### Build your automatic feeder

As already mentioned, ZAF is designed to be built with only commercially-available parts. Additionally, ZAF doesn't require 3D printing, machining, difficult adjustments, or the operation of heavy machinery. Most of the tools we will use are commonly found in an academic lab or easely sou.

Assuming that you have received all the parts you can build ZAF in one day! Start with the hardware, then assemble the electronics and finally install the software. To a large extent the speed at which you can build ZAF will depend on your D.I.Y skills.

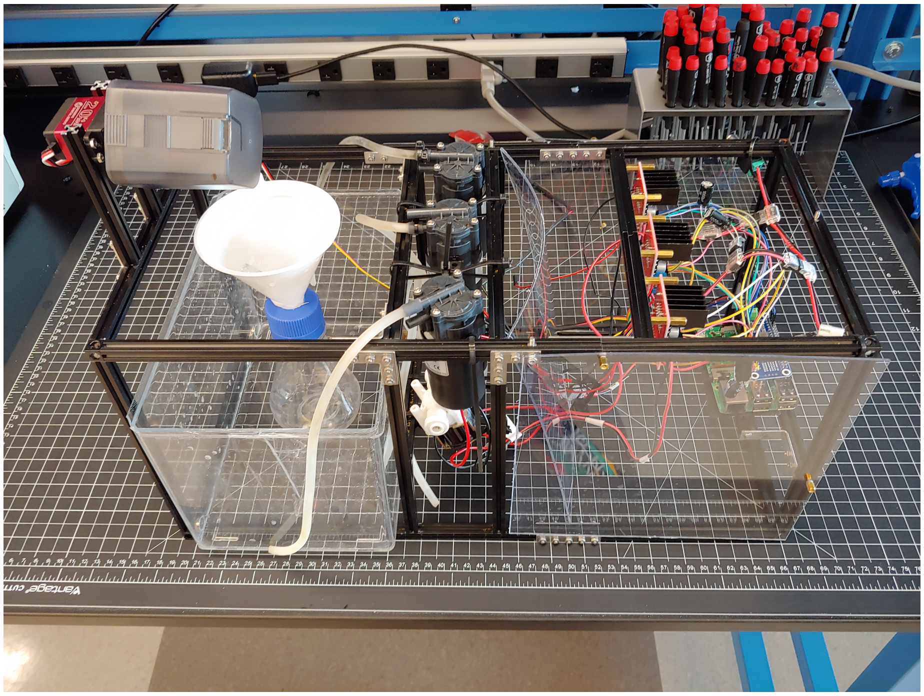

**Let's have fun with Makerbeam!**

To build our frame we used the Makerbeam starter kit. We provide detailed descriptions to assemble the ZAF frame, but you can easily adjust the sizing depending the number of pumps you need for your aquatic rack or facility.

Here are some basics, the frame must contain: 1. A frame compartment to hold the electronics: the Raspberry Pi, motor driver and screen. 2. A frame compartment to hold the pumps and valve. 3. A frame compartment to hold the water container where food and water are mixed and that holds the servo motor that rotates the food container.

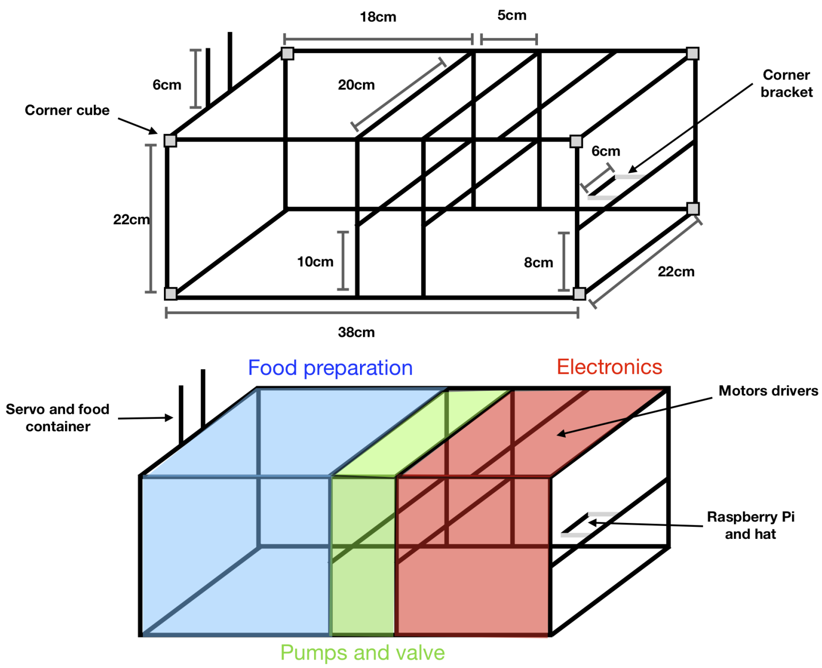


The frame design is important because it will be the “body” of your feeder. The Makerbeam corner cubes must be at the corners of your frame. Additionally you must anticipate the placement of the Makerbeam spacer and the Makerbeam square headed bolts, to support the electronics (*i.e* servo, motors drivers, etc. . . ) during the construction of your device structure.

There are many ways to use the Makerbeam steel brackets to construct your ZAF. In the following figure we give you an example of how to use the steel brackets to fortify the structure and build a custom support frame for the electronics. Do not hesitate to you use them intensively to get a robust structure.

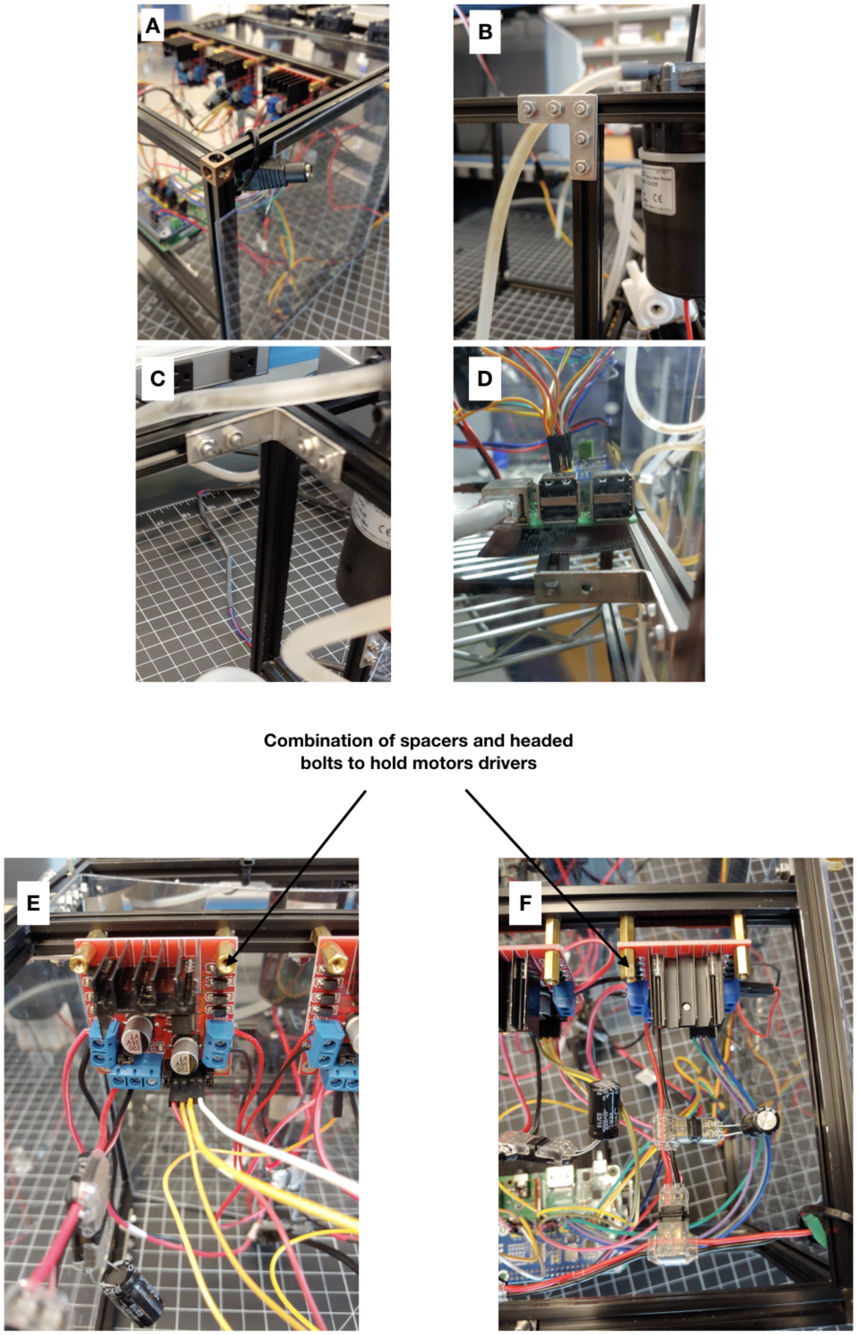


- A. Corner of our ZAF frame with a corner cube.
- B & C. Example of how to use the Makerbeam steel brackets to reinforce

the structure.

- D. We used the Makerbeam corner steel brackets and a small Makerbeam bar to create a support for the electronics component.
- E & F. Spacer used to hold Motors drivers

Now it is time to have fun and enjoy playing with the Makerbeam to build your own ZAF, a *Proust Madeleine* moment, appealing to your childhood memories playing Lego.

##### Ready for the food distribution?

To distribute the food that will be mixed with water and subsequently distributed to the fish tanks, we hijacked a food container obtained from a commercially available fish feeder and combined it with a Servo Motor. To attach the food container on the servo we used magnets to anchor it. Magnets allow the user to easily remove the food container for easy food refilling (Loic loves magnets and wants us to use magnets everywhere). By combining the servo and a container with a slider opening you can adjust the desired amount of food. Indeed, we can adjust the numbers of servo rotations and tune the container opening with the software. Each user can adjust these parameters to find out the right amount of food for their fish facility.

- Glued magnets on the food container and on the servo to create a magnetic anchor between the two parts. Use very strong glue.
- Be cautious about the magnet orientation, the food container opening must be oriented toward the top when placed on the servo.
- Fixed the servo on the robot body using Makerbeam headed bolt
- Done!

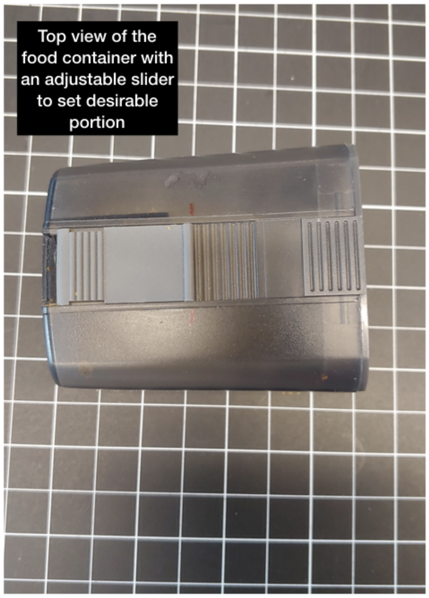
Top view of the food container with an adjustable slider to set desirable portion

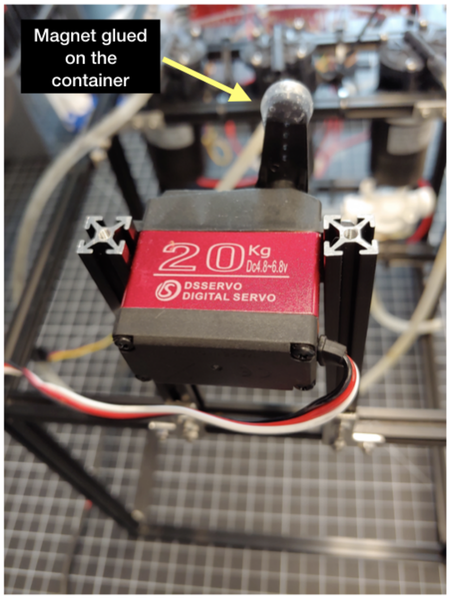
Magnet glued on the container
20Kg
Dc4.8~6.8v
DSSERVO
DIGITAL SERVO

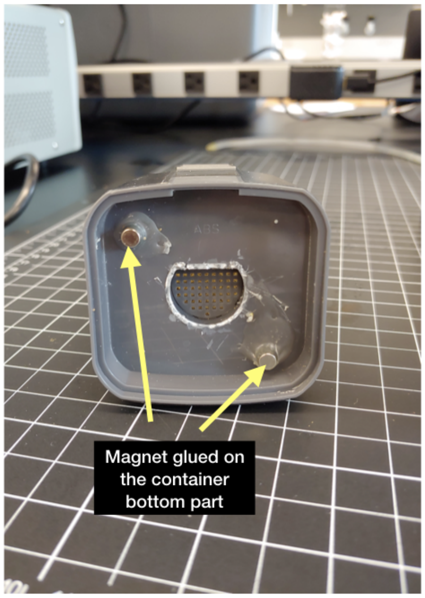
Magnet glued on the container bottom part

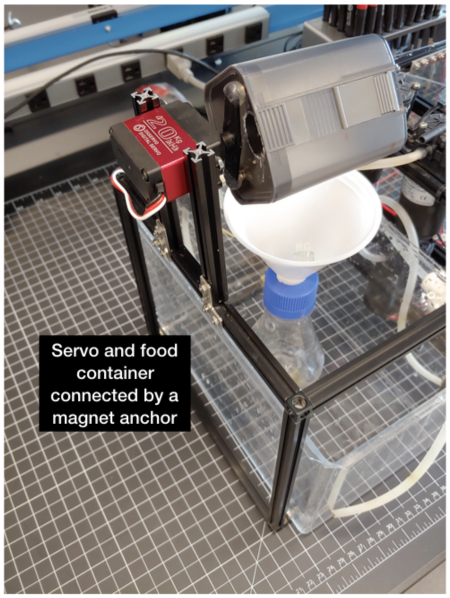
Servo and food container connected by a magnet anchor

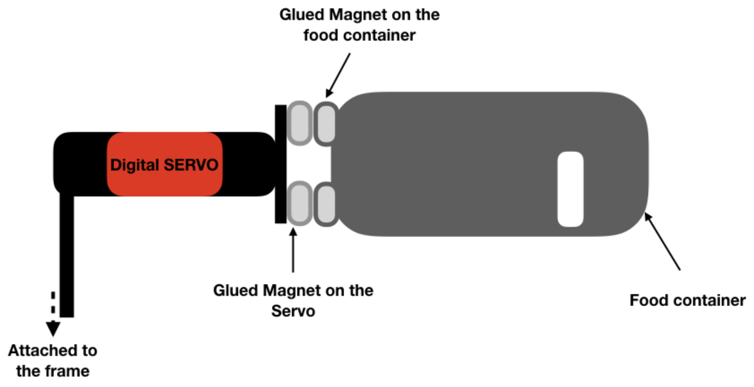


**It's time to setup the pumps!**

The pumps are the key elements to distribute the food and water mixture to the tanks. We decided to use micro gear pumps, that are self-priming, safe for food, and can move up to 2 liters per minute. We added a solenoid valve downstream of the "water in" pump to avoid water flow from the reservoir tank back into the ZAF device.

ZAF is designed to be modular, and therefore the number of pumps can be modified. In our original design, presented here, we have two pumps, and each pump can deliver food to 8 tanks. So just by adding pumps the user can scale up ZAF for more tanks. In this section, we will describe how to position the pumps and the valves in ZAF. Please refer to the following schema for where to place them and attach with nylon cables ties.

In the presented system we have one valve and four pumps:

- Pump food 1 & Pump food 2 –> Two pump for the water and food distribution to the fish tanks (feeding up to 16 tanks).
- Water in –> One pump to bring the water.
- Mixing –> One air pump to stir the food in the water.
- Air pump –> to mix the food and water.

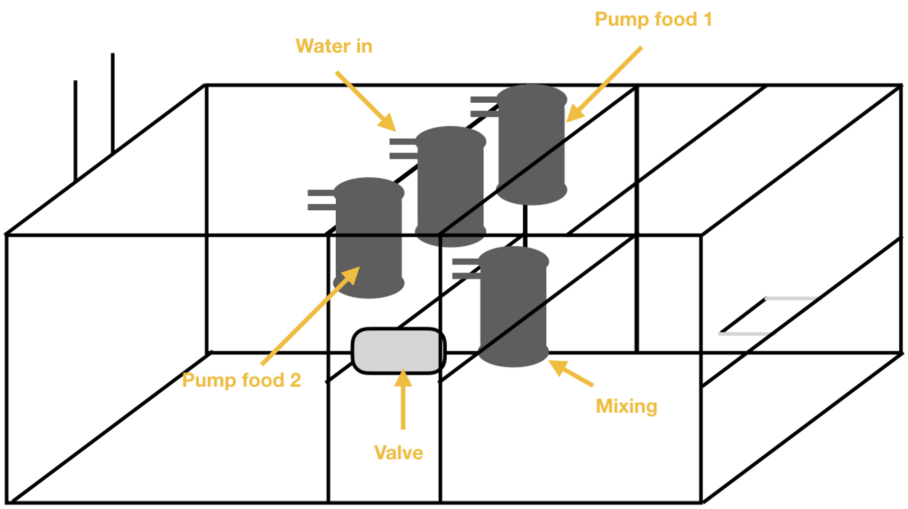


- A. Model of the micro gears pump used, with the in and out pipe.
- B. View of the three top pumps attached with nylon cable tie.
- C. Bottom pump and valve.

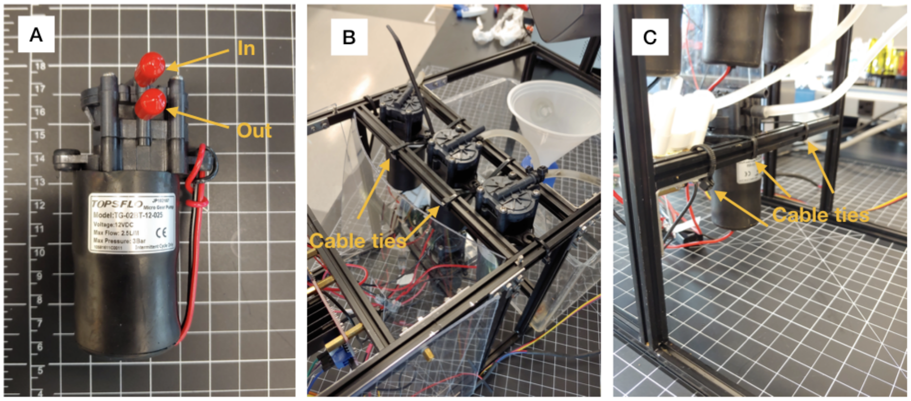


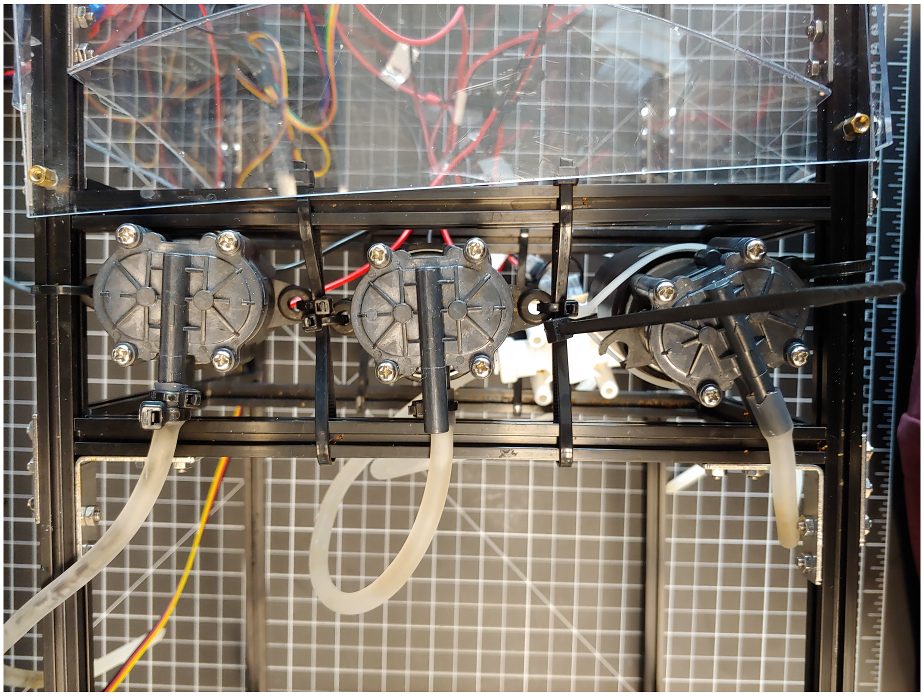

Top view of the pump area with the nylon cable tie.

Note: Readers who would like replicate ZAF might have hard time finding the exactly same brand and model pumps and valves we've used. Readers can use different brand/model pumps and valves with same technical specifications with what we've have used. We've chosen to use pretty generic pumps and valves, hence, it should be easy to find replacements if needed.

##### Time to prepare the 'kitchen area'

To mix the the food and the water we adapted a plastic 200 ml lab flask by drilling three holes on the top and added a funnel. The purpose is to insert the tubes connected to the pump: one for the tube coming from the mixing pump, and two connected to the pumps delivering the food to the tanks. Holes were drilled using a Dremel 3000.

We used the small funnel from this kit. The funnel is adapted on the lab shaker flask by inserting the output inside the cap of the aforementioned previously drilled flask. To maintain the tube inside the funnel we placed a tube holder suction clips.

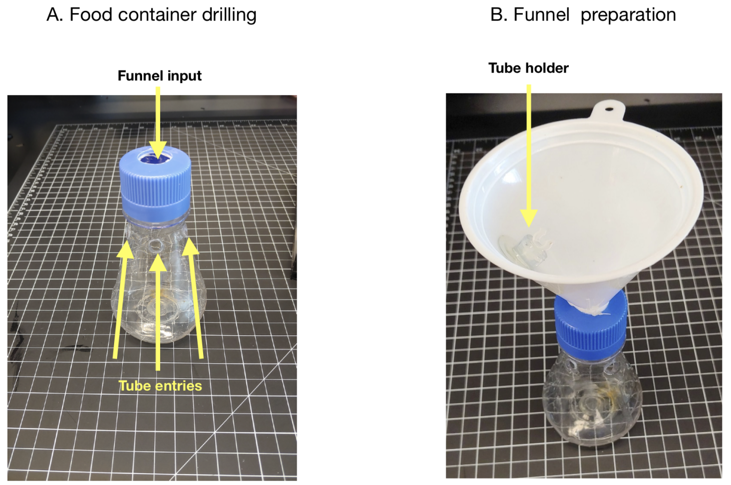


To prevent splashing and mitigate accidental leaks, the flask goes inside a water containment tank. We used a magnet anchor to maintain correctly the water container within the robot's frame. We glued two magnets on ZAF body and two magnets on the tank (be careful with the magnet's orientation!). Same glue used for the servo will work perfectly.

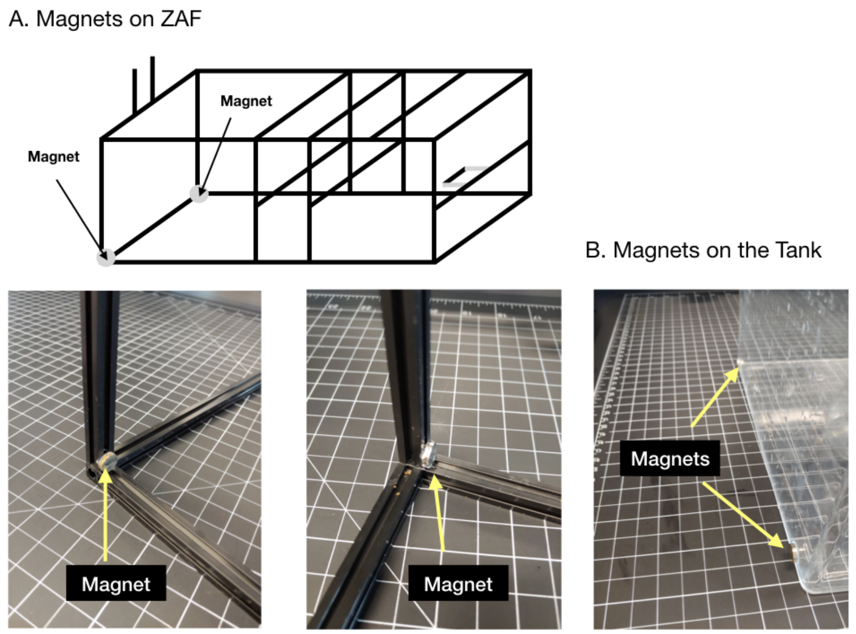


**Safety first!**

ZAF (and ZAF+) must run with minimal human supervision. The main failure mode is leaking due to splashing, or due to disconnected or clogged tubbing. Therefore, we added a security system to avoid accidental flooding. The safety container is the first line of defense. However, in the case of serious leaks, water will eventually accumulate. To prevent an excessive amount of water in the safety container, we installed a water sensor coupled to a water submersible pump. In the case of a leak, the sensor detects water in the tank to activate the pump to remove the excess water.

Furthermore, in order to avoid electronic failures we embedded the top part of the sensor in silicone. We anchored the pump using the magnet trick described before to attach the the water container tank to the ZAF body and the servo and food container. The sensor is attached to the tanks using a tube holder suction clip like the one used in the funnel of the food preparation flask.

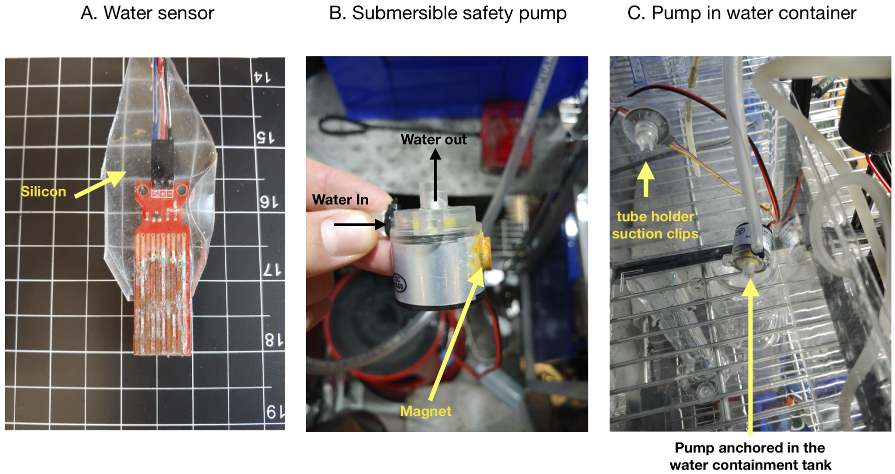


#### Tubing for food distribution

Tubing delivers food to the tanks and is thus an essential element of the device. You must carefully design and assemble the tubing elements (i.e, tubes, splitters, connectors) to prevent any problems during food distribution. Here we provide a general framework for ZAF tubing, as well as detailed instructions on how to assemble the splitter panel. The splitter panel is a key elements to separate the flow of food and water and deliver an equal amount of food to the different tanks.

#### Tubing

For ZAF we used silicone based tubes, make sure to use tubes that are safe for the fish (food grade!) and that are rigid enough to prevent clogging. We used two types of tubes: type 1 for connecting pumps valve and food flask inside ZAF, and another: type 2 to hook up the tanks with the splitter panel.

In this schematic graph we indicate how to connect the different pumps and valves together, and what type of tubing is used.

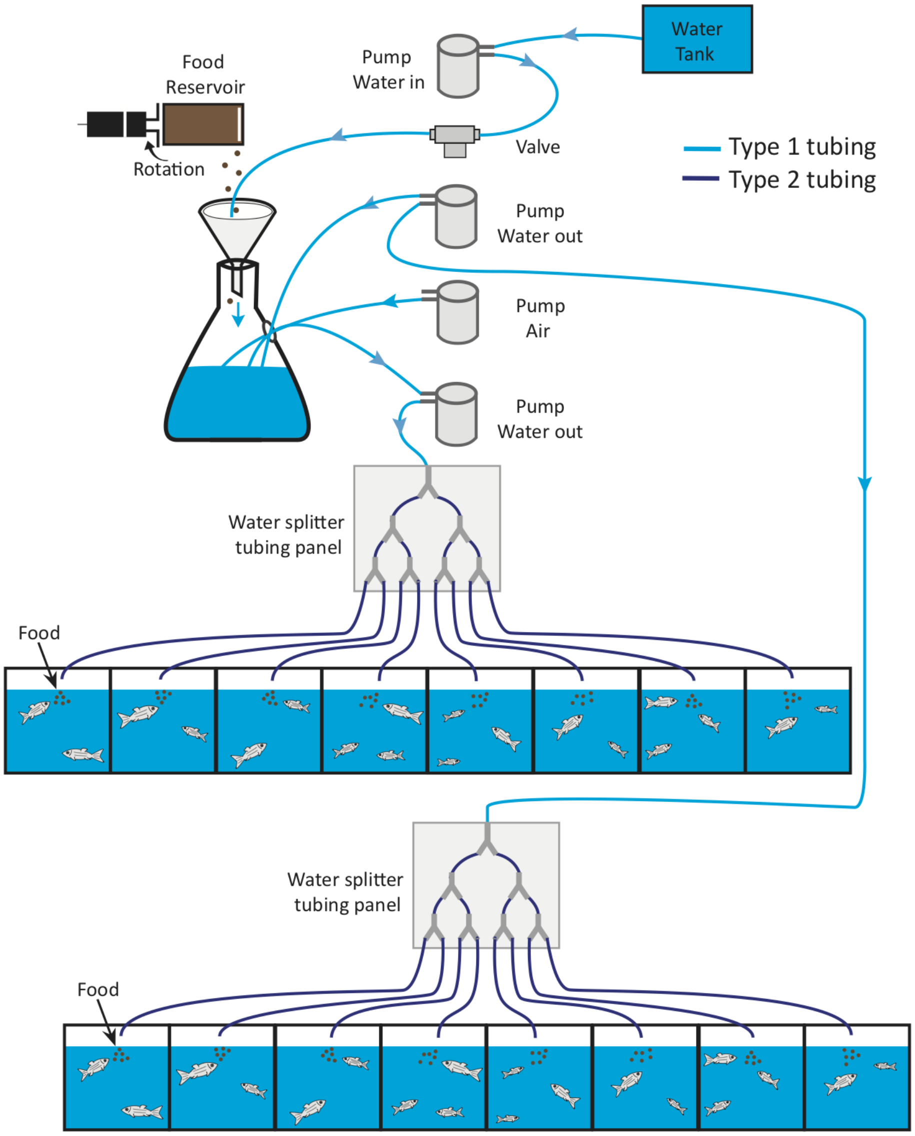


Keep in mind that you will need to keep all tube lengths equal, even for different tanks. Indeed, we want to ensure that the tubing resistance to water flow is equal across all lines so as to guarantee the equal distribution of food across all tanks. To avoid cable overcrowding the cable tie can be used to organise the tubes.

##### The water separator tubing panel

These are the “tentacles” of your robot that will distribute an equal amount of food to each tank. In the described design, one separator panel (connected to a single pump) delivers food to eight tanks. We decided to limit our panel to eight tanks but with ZAF it is all about modularity. The users can try a panel with 12 tanks! However, scaling up to many more tanks runs the risk of clogging because of reduced pressure – so for more tanks we recommend ZAF+.

The panel support is made of polycarbonate plastic sheet with dividers fixed using tube holder suction clips. We used two Y-dividers, the type 1 tubes for the first coupling and then the type 2 tubes for the output connections. The middle T-connector are from this kit.

Overall, it is easy to assemble the panel following the indications on the following figure (left panel). On the right it is a picture of the panel inside our facility. Finally, we fastened the panel using velcro stripes.

We recommend the users to evaluate the aspect of the tubes regularly (water separator panel as well as system) and replace the tubes when needed.

**Remember: all tube lines, must all have the same lengths**

The water separator tubing panel

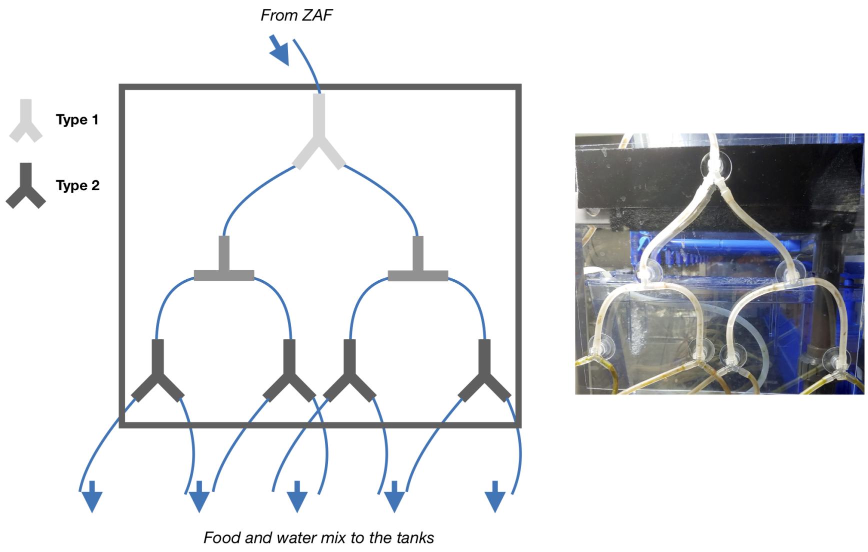

##### Time to take control of ZAF

At this point you should have managed to build ZAF's frame and the main hardware components. Let's now give the feeding device a nervous system. We will describe in this section how to connect the different components to control ZAF with the software we developed (described in the last section). The electronic core of the design is based on:

- A Raspberry Pi 3 B+ a credit card size computer.
- A Servo Hat Board to drive PWM outputs, like the pumps and valve.
- Motor Controller to control the DC motors (pumps and valve).

All the pumps and valves connected to the motor drivers are plugged on a 12V and 10A power supply converter. To help stabilise the voltage, and reduce voltage ripples due to the motors turning on and off, we connected 1000uF capacitors in parallel to the motor terminals. The Raspberry Pi, the servo hat and all the electronic connected to the servo hat are running with 5V through the Raspberry Pi power.

You can find a complete description of the electronic setup in the following figure:

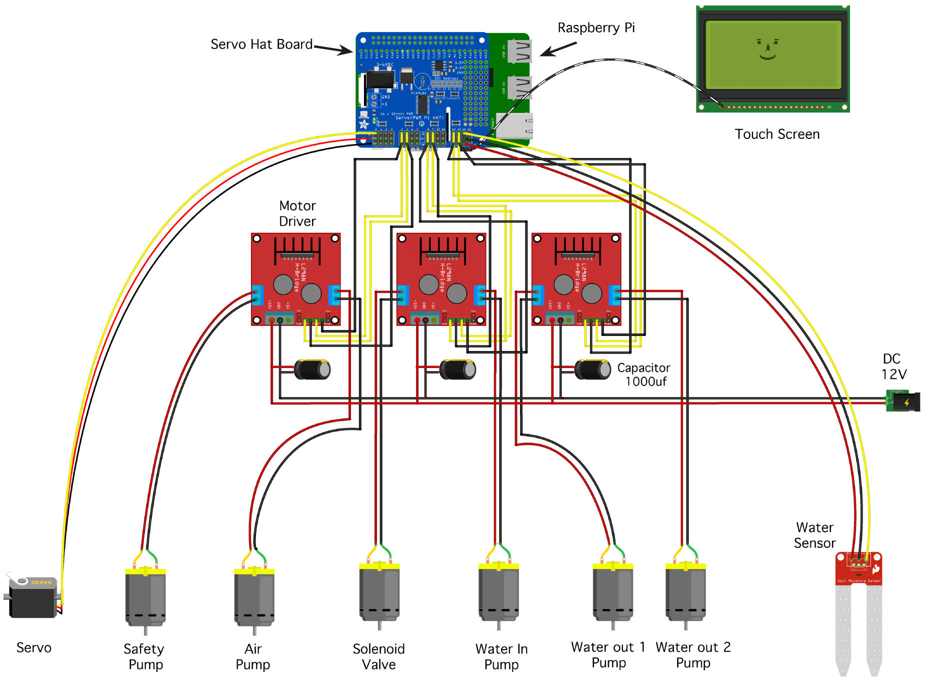


##### ZAF is solderless!

The first version of ZAF is 100% solderless. We used splice connectors, either T-type or I-type. These connectors are cheap, safe, and very easy to use! However, if you prefer, you can use soldering to connect all your wires and devices together.

Importantly, you should aim for a clean setup with well-connected cables and no loose wires. To further secure the connections, you can use 'electrical insulating tape'.

Overall, when placing your robot in the fish facility, make sure to place the electronics – and in particular the cables carrying 110 or 240V – far from water and at a safe distance away from splashing. In practice we found that elevating ZAF to the height of the highest tanks is a good strategy both for the flow of water and to avoid the risk of ZAF electronics being flooded.

Here we show how to use the splice connectors:

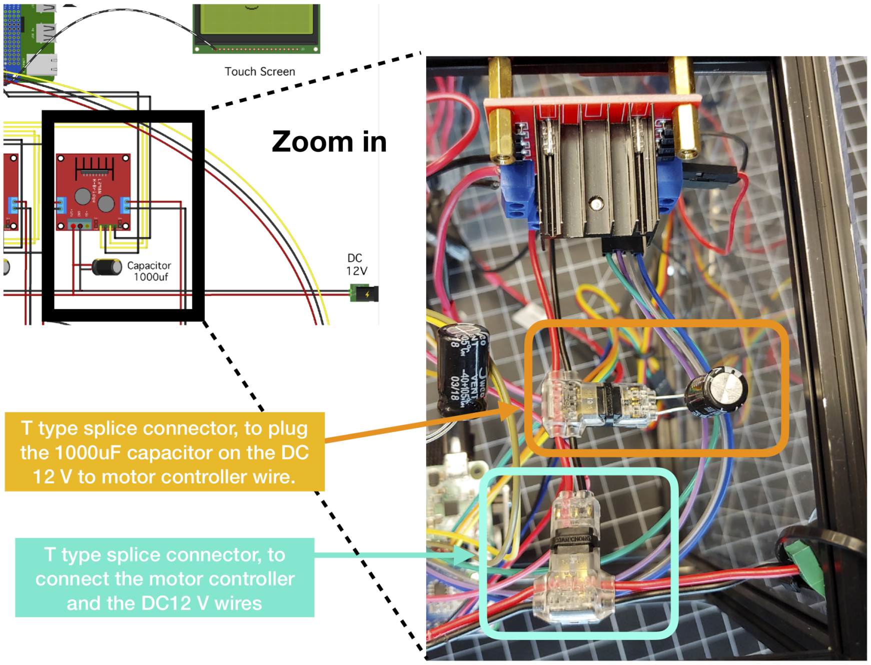


##### Motor Drivers the optimal pump controller

The motor drivers (L298N) use standard mosfet-based H-bridges to control the voltage polarity at its output connectors. The servo hat controls the motors drivers via 5V TTL lines and can modulate these using Pulse Width Modulation (PWM). For our purposes, this gives us the ability to precisely control our pumps and valve voltage.

Overall, connecting the different components is easy: one motor driver can be connected to two output devices, and each motor driver can be controlled by the servo hat board, for a theoretical maximum of 8 pumps or valves. Each motor driver is connected to a shared DC 12V 10 A power supply, with a 1000uF capacitor in parallel for each driver.

Exact details on how one motor is connected are described in the following figure:

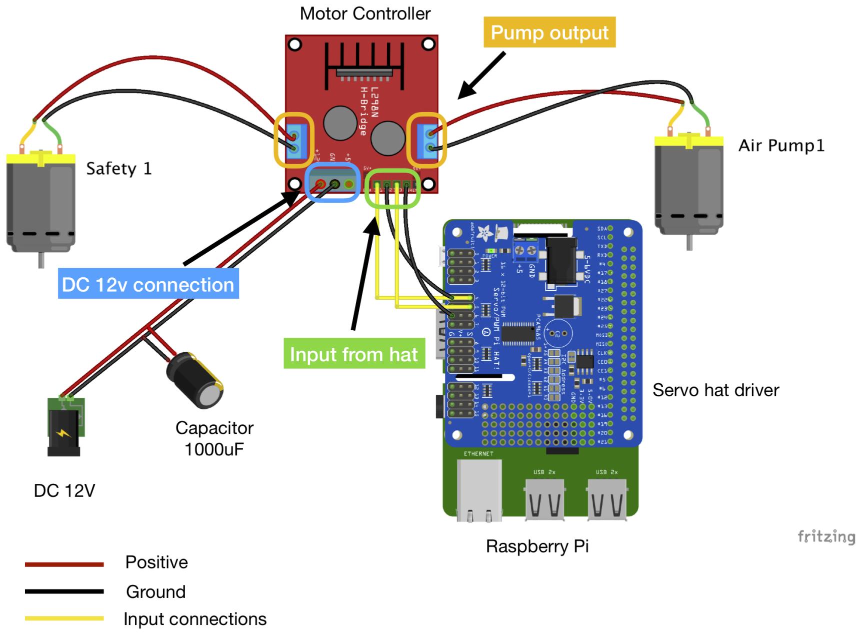


In the main diagram you can find all the specific connections needed. To summarise:

- One motor driver for the safety pump and air pump
- One motor driver for the solenoid valve and the water-in pump
- One motor driver to water out 1 and water out 2 pumps (the pump distributing the mix of food and water to the tanks)

**Reminder – ZAF is modular and extra motor drivers can be added to accommodate additional pumps!**

##### Time to plugin the micro computer and the servo hat.

For ZAF you need to use a Raspberry Pi 3 B+ microcomputer. We used a kit to get most parts needed to operate the Raspberry Pi – including the case. To attach the Raspberry to ZAF's frame we used velcro as described in the tubing section. To setup your Raspberry Pi please go here.

Next you will need to connect Raspberry Pi to the servo hat board. This board is called a 'hat' because it is attached on top of the Raspberry Pi and communicates with the Raspberry Pi through the GPIO connectors (here for a detailed explanation).

**Servo Hat connection.**

Your servo hat sends PWM signals to the motor controllers to control the pumps and valves. Other devices, such as the servo or safety pumps, can be directly controlled with 5V TTL lines directly from the servo hat.

Next, we describe the connections between the electronics devices and the servo hat board:

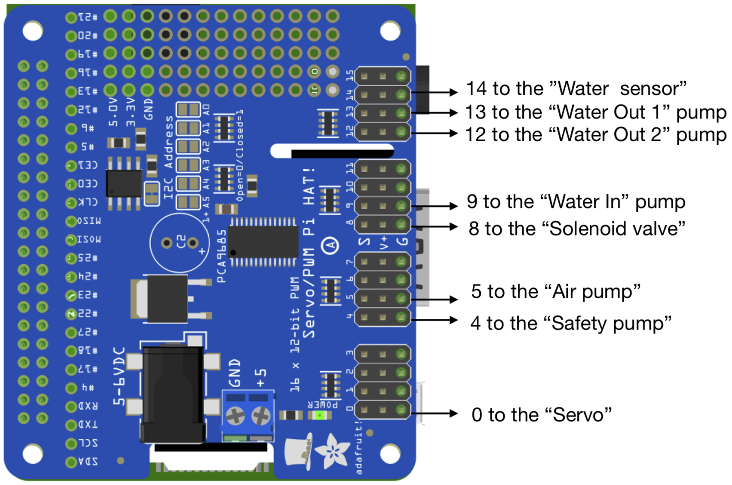


- For the wires coming from the Motor controller connect the Input 1 and 3 (IN1 & IN3) to the raw "S", for signal, on the Servo hat. And the Input 2 and 4 (IN2 & IN4)on the ground "G". You can check the details on the motor drivers page.
- For the direct connection (water sensor, servo and safety pump), connect directly voltage plus and minus and the signal "S" on the servo hat board.

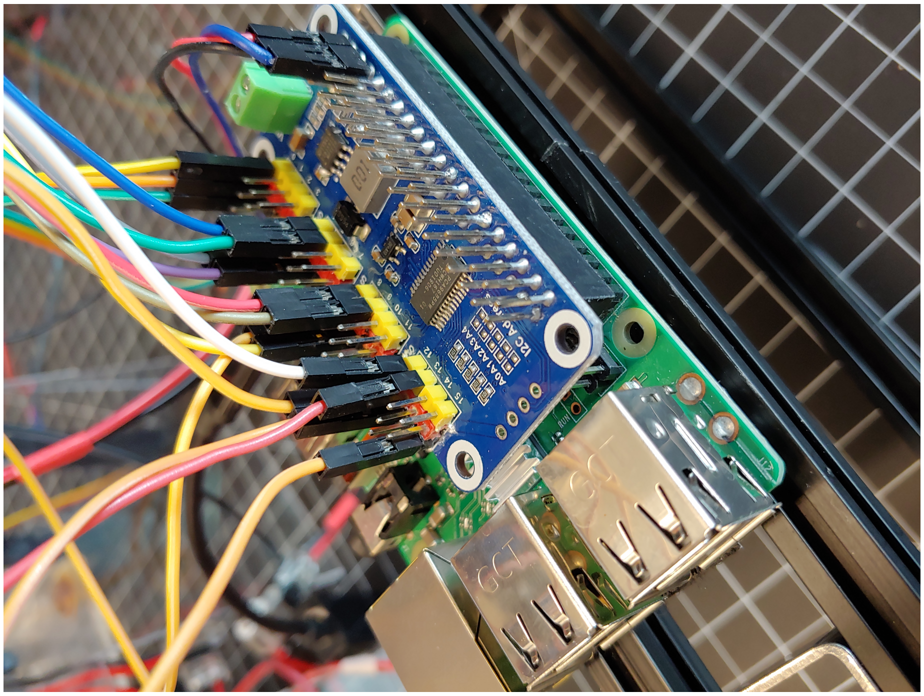

We built a module for the Raspberry Pi 3 B+ and the touch screen using Makerbeam. To set up your Raspberry Pi for easy control, we added a wireless keyboard with a touchpad to the system. Follow this instruction to set up your Raspberry Pi. The touch screen and the Raspberry are connected together with an HDMI cable and a USB/micro-USB cable. For details on the connections and how to fasten them to the Makerbeam structure, please refer to the following pictures.

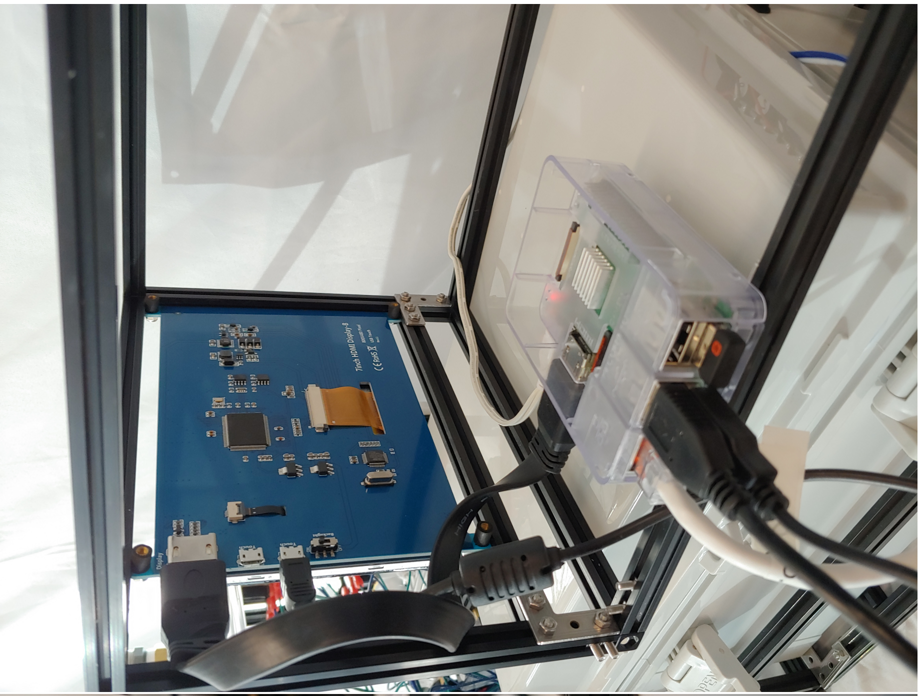

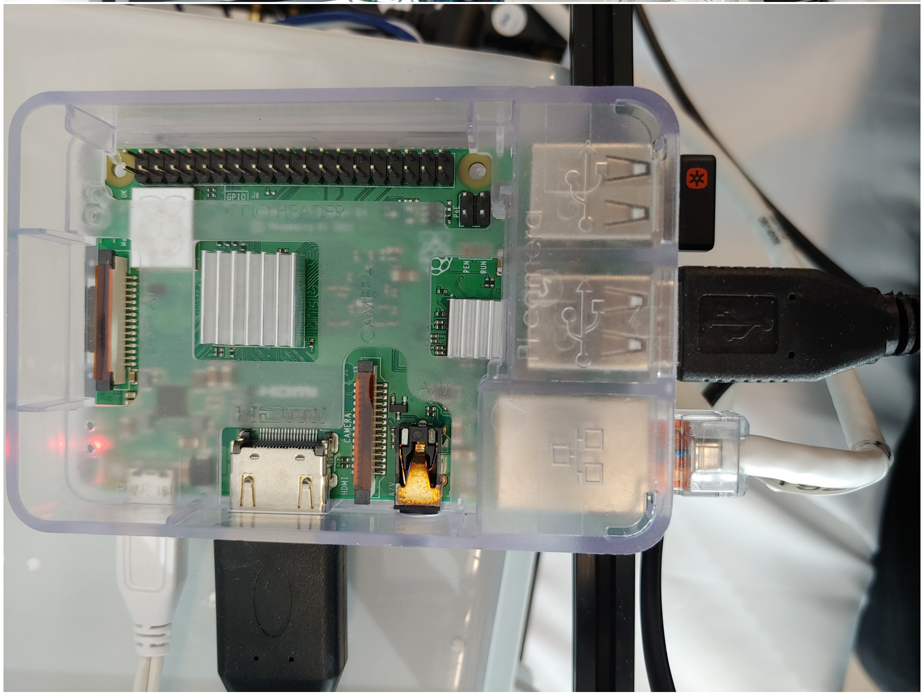

### Install and run the ZAF control software

The standard ZAF hardware is controlled by a very simple python-based program, below are the instructions for installation, usage and calibration. This software piece implemented and tested on Raspberry Pi with Python2.7.

- Installation
- How to use
- ZAF calibration # Installation

**Preparing the Raspberry Pi** ZAF code is written and tested on only Raspbian Buster >=10.

- First, we recommend you first have a look at the official documentation.
- To get started first download the Raspbian OS and burn it in your microSD card. We recommend doing this with the help of Raspberry Pi Imager which makes this process painless.
- Then you need to get command line access to your Raspberry Pi. You can connect it to a local network via ethernet or wifi. Once connected to the network, you can access a remote terminal using the ssh protocol. A simpler approach consists in connecting a monitor and keyboard to your Raspberry Pi and opening the terminal.
- After having access to Raspberry Pi terminal, follow the commands listed below one-by-one:

You can basically get the ZAF source code by cloning our repository into your raspberry pi and start using.

You can get ZAF source code by:

```
pip install click
git clone https://github.com/royerlab/ZAF.git
```

- Congratulations! You successfully installed ZAF software.

### How to use

#### Command Line Interface(CLI)

ZAF software provides a small command line interface(CLI). With the help of CLI one can run the ZAF with

```
python -m python.zaf.cli run
```

Other commands are not implemented yet. There two other(`last5`, `last50`) dummy commands not doing anything particular at the moment.

##### Scheduling Feeding

It is very useful to be able to schedule feeding sessions. ZAF software has no builtin feature to have scheduled feeding sessions but this can be achieved by using crontab on Raspbian OS.

To create new crontab tasks or edit existing tasks one can do:

```
crontab -e
```

To list the existing crontab tasks you can do:

```
crontab -l
```

#### How to scale up ZAF?

ZAF design is about modularity so without major modifications you can easily implement more feeding outputs.

The first steps is to multiply the hardware. Adding extra pump and therefore all the complementary electronics, connection of a motor driver to the servo Hat bard, and tubing can increase ZAF capacities. You can add 8 extra LN298 to one servo hat and each of them can be connected to 2 pumps. Thus, one ZAF can distribute foods up to 80 tanks!! You can also simply connect bigger pumps (similar to the one use in ZAF+) and then multiply the number of output in the water separator panels.

You will also need to adapt the code.

- If you like to add a new pump, you need to first register the new pump in `Ctx` class we have in `python.zaf.ctx` submodule.
- Then, you can go to `python.zaf.fishfeed` submodule and use the new pump at any point of the code you like.‘
- For instance if you added a new `water_in` pump, you can adapt your code to use two `water_in` pumps as shown below:

```
Ctx.pwm.setPWM(Ctx.water_in, 0, 4095)
Ctx.pwm.setPWM(Ctx.water_in2, 0, 4095)
sleep(0.5)
Ctx.pwm.setPWM(Ctx.water_in, 0, 0)
Ctx.pwm.setPWM(Ctx.water_in2, 0, 0)
```

If you have have further questions about scaling up ZAF feel free to contact us by opening an issue on our repository. ## Necessary Equipments

The table is indicating the basic equipments we used to build our ZAF. Most of the equipments used is generic and can be replaced by components with similar specs. We added hyperlink on the GitHub wiki to help the user buying the appropriate materials. The only components that cannot be easily exchanged is Raspberry PI, but that is not an issue as these are very easily sourced components.

| Components | Parts name | Supplier | Article Number | Number | Unite Price (USD) | Total Price (USD) |
|---|---|---|---|---|---|---|
| Frame | Makerbeam Starter Kit | Makerbeam | 103318 | 1 | 112.25 | 112.25 |
| Servo & Food container | Eheim Automatic feeding unit | Eheim | NA | 1 | 23.47 | 23.47 |
| | Digital Servo | N/A | DS318 | 1 | 16.66 | 16.66 |
| Pumps & Valve | Magnets | Dymag | 3MM-mix | 1 | 14.99 | 14.99 |
| | Pumps | Trossen Robotics | TI-TG-02B-DC12B | 4 | 24.95 | 99.8 |
| | Valve DC 12V ¼” | Digiten | DC 12V ¼” | 1 | 7.49 | 7.49 |
| Food Mixing | Cable ties | McMaster-Carr | 80005k2 | 1 | 25.05 | 25.05 |
| | Tube Holder | Asayu | N/A | 1 | 5.99 | 5.99 |
| Safety | Funnel | Karzone | N/A | 1 | 6.99 | 6.99 |
| | Water sensor | DAOKI | TS-VS-292-CA | 1 | 20 | 20 |
| | Minipump | Walfront | 12V DC 6W | 1 | 15.49 | 15.49 |
| Tubing | Type 1 tubing 3/16” | CNZ | N/A | 1 | 5.99 | 5.99 |
| | Type 2 tubing 4mmID, 6mm OD | NACX | N/A | 2 | 22.88 | 45.76 |
| | Polycarbonate plastic sheet | Robosource | N/A | 1 | 24.99 | 24.99 |
| | 40 aquarium connectors | Asayu | N/A | 1 | 11.99 | 11.99 |
| | Velcro strips | Velcro | S-5751 | 1 | 37 | 37 |
| Electronics | Wire connectors T-type | Biantie La | N/A | 2 | 9.98 | 19.76 |
| | Wire connectors I-Type | Biantie La | N/A | 2 | 9.99 | 19.98 |
| | Motor Drivers | Qunqi | L298N | 2 | 8.69 | 17.38 |
| | Canakit Raspberry Pi 3B+ | Canakit | N/A | 1 | 69.99 | 69.99 |
| | Servo Hat 16 Channel | Adafruit | 2327 | 1 | 17.5 | 17.5 |
| | LCD Touch Screen 1024X600 | Longruner | LSC7B | 1 | 56.99 | 56.99 |
| | | | | | **Total Price** | **675.10$** |

##### It is time to build the complete automatic solution for fish feeding

ZAF+ is an extension of ZAF. The two devices share many similarities but ZAF+ is an improved version of ZAF that allows for precise food distribution thanks to valves placed upstream of each tank. While ZAF+ is more complicated to build, we have kept the original ZAF principle: it is designed to be as simple as possible, does not require complex machining or 3D printing, and all components are commercially available and can be assembled using common tools.

Here we describe the construction of ZAF+ scaled to feed up to 30 tanks – a typical average number of tanks found in most standalone zebrafish racks. We provide here a straightforward hardware and electronics assembly manual for building ZAF+. ZAF+ is modular and can be adapted to the user's facility needs and scaled accordingly. Additionally, the number of valves can be adjusted to provide food to a larger number of tanks, the frame can be optimized to the fish facility layout, etc… The customization possibilities are numerous!

#### Picture of the ZAF+ built for the wiki

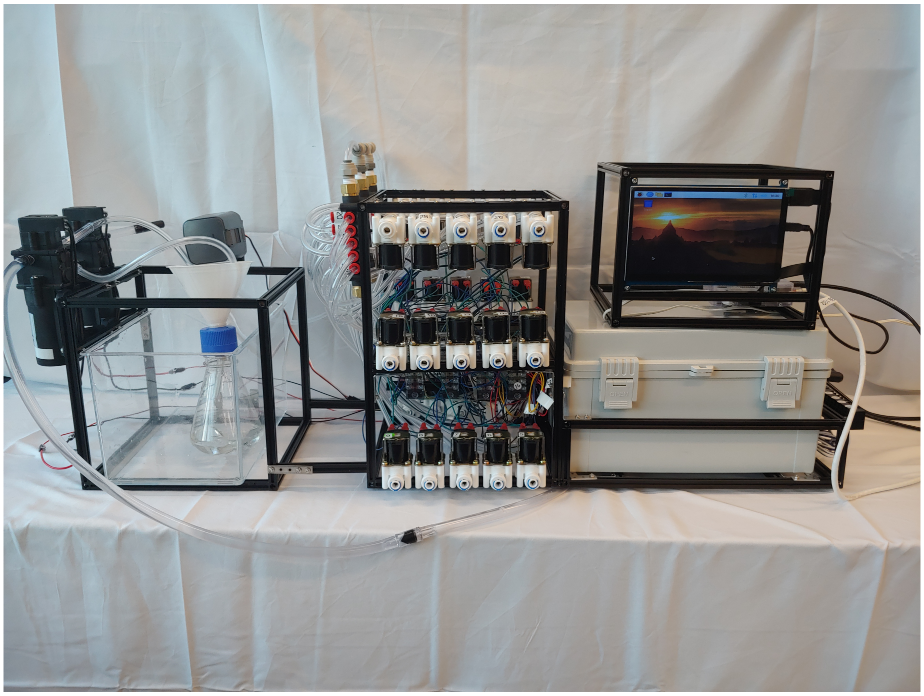

Figure 2: ZAF+

##### Modularity adapted to the different ZAF+ components

Similarly to ZAF, you will need a Makerbeam starter kit. However, we recommend that you buy more Makerbeam kits since the ZAF+ structure is more elaborate and needs to support more components. Of course, any similar prototyping beam-based system could work too, and we welcome suggestions for alternatives. For the model presented herein you will need two packs of 200mm, two packs of 300mm and a bag of corner cubes. Of course, this list is not exhaustive and can be readily adapted to your specific design needs. Do not hesitate to navigate on the Makerbeam website (or alternative) to find the ideal components for your ZAF+ project.

The ZAF+ frame is divided into four modules that can be attached together, each responsible for a different function during automated feeding: 1. Raspberry Pi computer and touch screen. 2. Electronics for motor and valve control. 3. Valves and tubing assembly. 4. Food preparation and pumps.

There are many different ways for you to build ZAF+ using the Makerbeam components. In any case, the design must integrate the four modules described above. We apply the same principal as for ZAF: the structure must be solid and integrate in its original design the location of the different components (e.g valves, pumps, water manifold, etc...). Check out the ZAF frame page for some examples, such as structure reinforcement with steel brackets and also how we built electronics support using Makerbeam accessories. To optimize the space and the ergonomics of ZAF+, we directly drilled in the 20mm beams to be able to use screws at any position along the beam. We used the drilled holes to fix the valves and the manifold on the Makerbeam 200mm and to attach the pumps tightly to the structure.

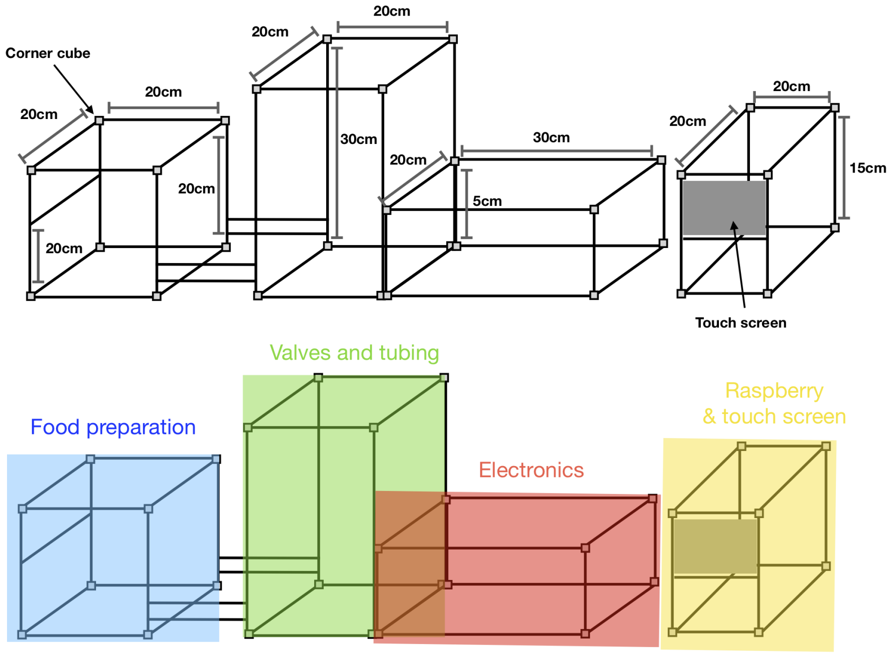


The "kitchen" for ZAF+ is built similarly as for ZAF here: it consists in a servo and a food preparation container. We recommend to use a bigger flat-bottomed flask for water and food mixing. We decided to use a larger container for ZAF+ because we use more water.

We used a servo motor to rotate the food container and distribute a certain amount of dry food pellets to each tank. The system used for ZAF+ is the same described for the ZAF here. Note - we also tested the system with fresh Artemia nauplii and it works well. If your fish diet includes *Artemia* nauplii, you can use ZAF+ for the food delivery, but you will have to prepare the food daily. For operation with less supervision we recommend using ZAF with dry food.

ZAF+ can control the quantity of food per tank, which is defined in the software using the Graphic User Interface (GUI). We implemented four food quantities: tiny, small, medium, large. Each quantity can be further defined during the GUI calibration depending on the user's needs. To adjust the amount of food we change the rotation of the servo, and therefore the amount of food delivered to an individual tank.

Based on our dry food seller's recommendations (we used the Gemma 500 from Skretting), adult fish need around 15mg of food per day. Therefore, based on this number we calibrated our ZAF+ for:

- Tiny - for 2 to 4 fish approx. 100mg of food delivery
- Small - for 5 to 8 approx. 200mg of food delivery
- Medium - for 9 to 14 approx. 350mg of food delivery
- Large - 15 up to 20 approx. 500mg of food delivery

**Scaling up and more precise food distribution.**

We control the distribution of the food mixture using 12V water solenoid valves upstream of each tank to deliver a precise amount of food into each tank. The software allows the user to control the individual valve upstream of a specific tank during the food distribution, therefore controlling food quantity.

The feeding procedure is as follows: a user specified amount of food proportional to the numbers of fish per tank is dropped, then the valve upstream for a given tank is opened, and finally the associated pump is turned on flowing a mix of water and food into the tank.

Importantly, in this version of ZAF+, each tank requires its own 12V water solenoid valves. In our case we wanted to provide food for 30 tanks, therefore we needed 30 water solenoid valves (food grade). We also decided to upgrade the pumps (compared to ZAF design), with diaphragm self priming pump with a flow up to 4.5L/mn. One pump is needed to bring the water to the system and another pump for food delivery. To avoid water flowing in reverse, we added check valves upstream and downstream of the pumps.

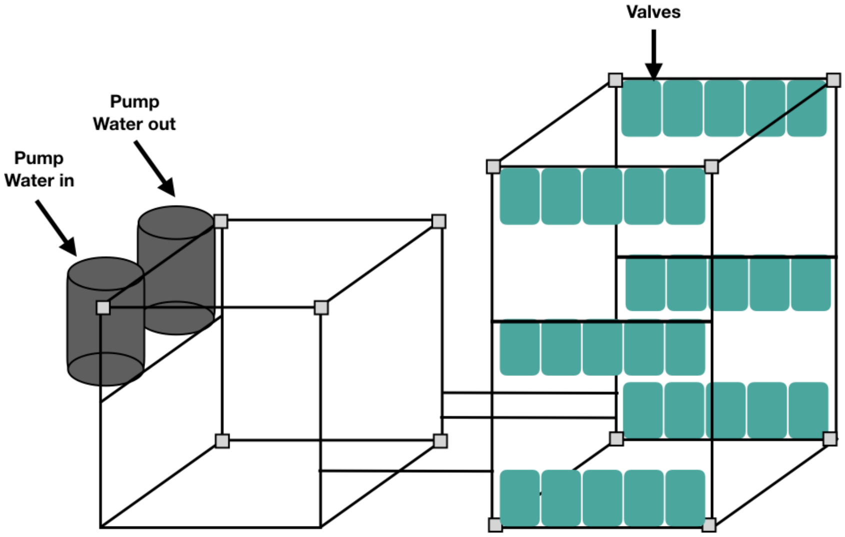


- Schematic representation of the pumps and valves assembly in the food preparation and valves and tubing frames respectively.

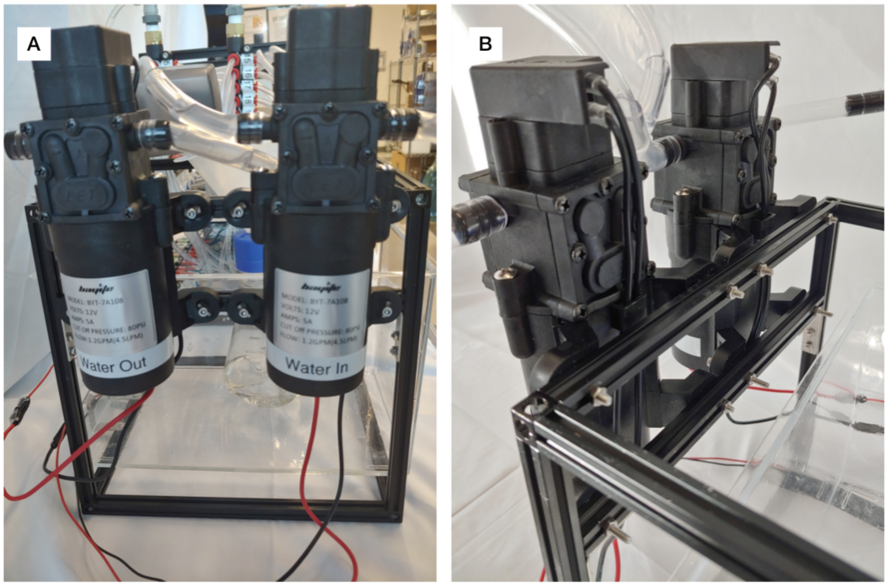


- A&B. Mounting of the pumps on the frame

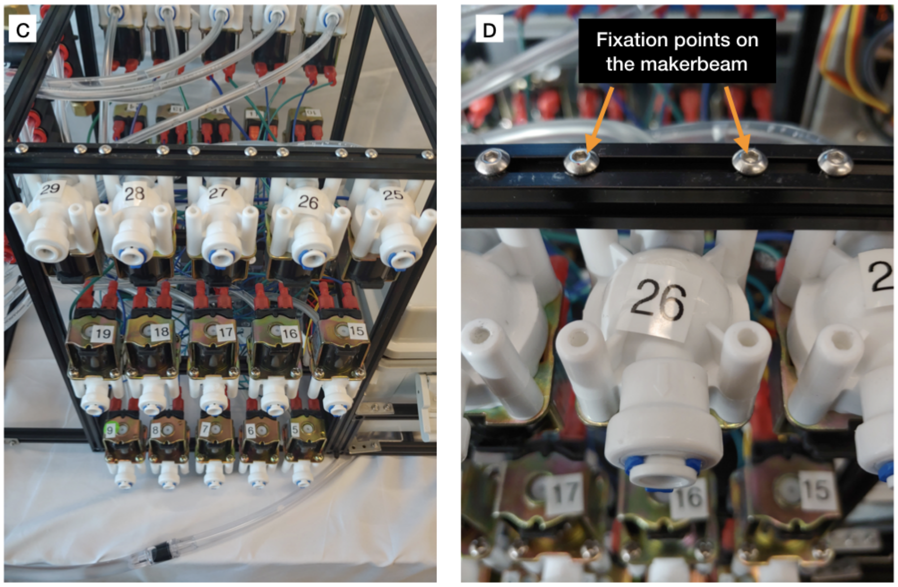

- C. Valves mounted in series of 5 on rows
- D. Fixation of an individual valve on a drilled Makerbeam 200mm

Note: Readers who would like replicate ZAF might have hard time finding the exactly same brand and model pumps and valves we've used. Readers can use different brand/model pumps and valves with same technical specifications with what we've have used. We've chosen to use pretty generic pumps and valves, hence, it should be easy to find replacements if needed.

We used a similar approach in both ZAF+ and ZAF and employed a safety container and a security pump. You can follow the assembly procedure here.

For safe automatic feeding without any human supervision, we installed several wireless cameras in our facility to remotely watch the feeding process. We also monitor for any system leaks using wireless water rope sensors, Monit. Altogether, combined with ZAF for the feeding, these solutions allow a safe remote maintenance of aquatic facility.

##### Pumps, manifolds and valves connections

The tubing is an essential component of ZAF+, just as for ZAF, but with a twist. In ZAF+ the totality of the food liquid mix is distributed to each tank. In order to connect the pump to multiple valves (30, in our case) we used manifolds that split the water flow into several tubes. Therefore, the food is distributed to one tank by opening the valve downstream of the pump and upstream of the tank to be fed.

The outside diameter of the *pump tubing* (3/8" from McMaster 5233k65) and the *valves tubing* (1/4" McMaster, 5233K52) is different so we used some push-to-connect straight reducers (McMaster, 5779K355) to connect the two differently-sized tubes. In ZAF+ we used 3 manifolds (McMaster, 52045K216), to circulate the water to up to 30 tanks using only one distribution pump. The first Manifolds are connected using t-shape push to connect adaptor (McMaster, 51055K27) and the last one with a right angle push-to-connect adapter (McMaster, 51055K195). We used an empty fish tank of the fish facility as *the water reservoir tank* to supply water to ZAF+.

Note - this time we used PVC based tubing because they are cost effective and have good specifications for ZAF+ food delivery.

The following schematic diagram represents the tubing connections between the different ZAF+ components (e.g, pumps, valves, food preparation containers, fish tanks etc…).

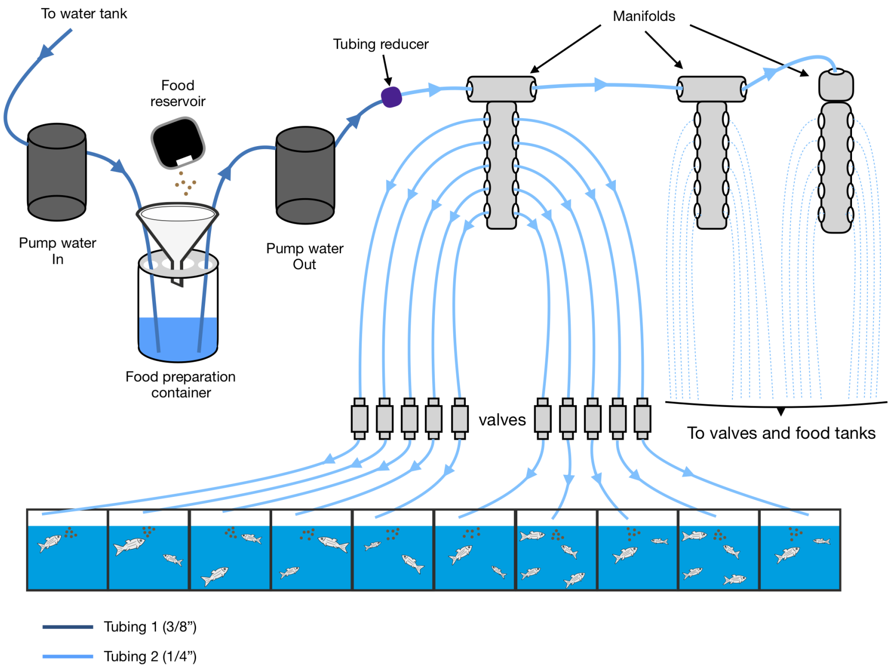
To water tank
Food reservoir
Tubing reducer
Manifolds
Pump water In
Pump water Out
Food preparation container
valves
To valves and food tanks
Tubing 1 (3/8”)
Tubing 2 (1/4”)

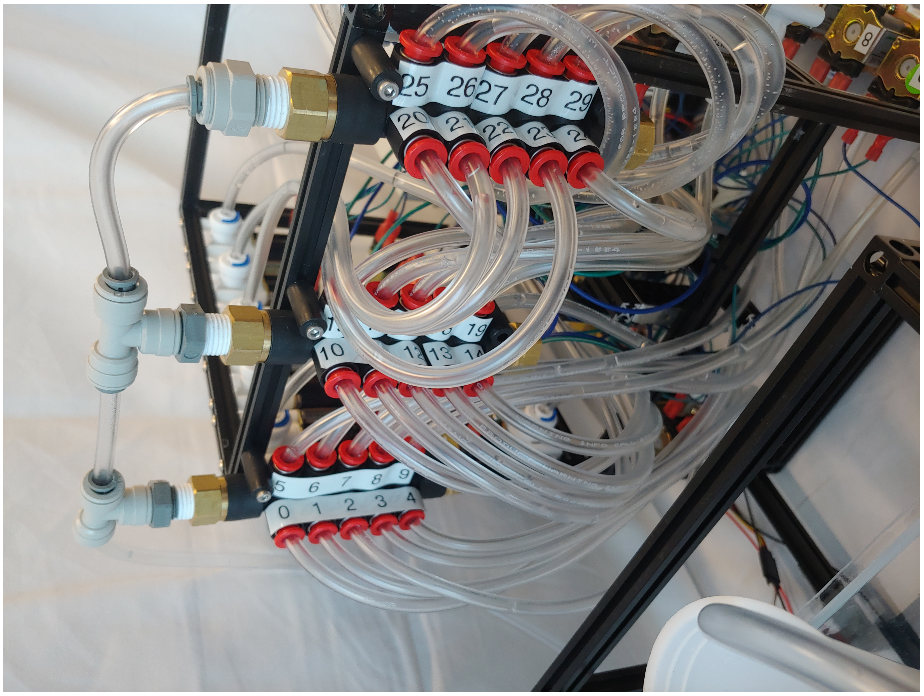
25 2627 28 29
10
5 6 7 8 9
0 1 2 3 4

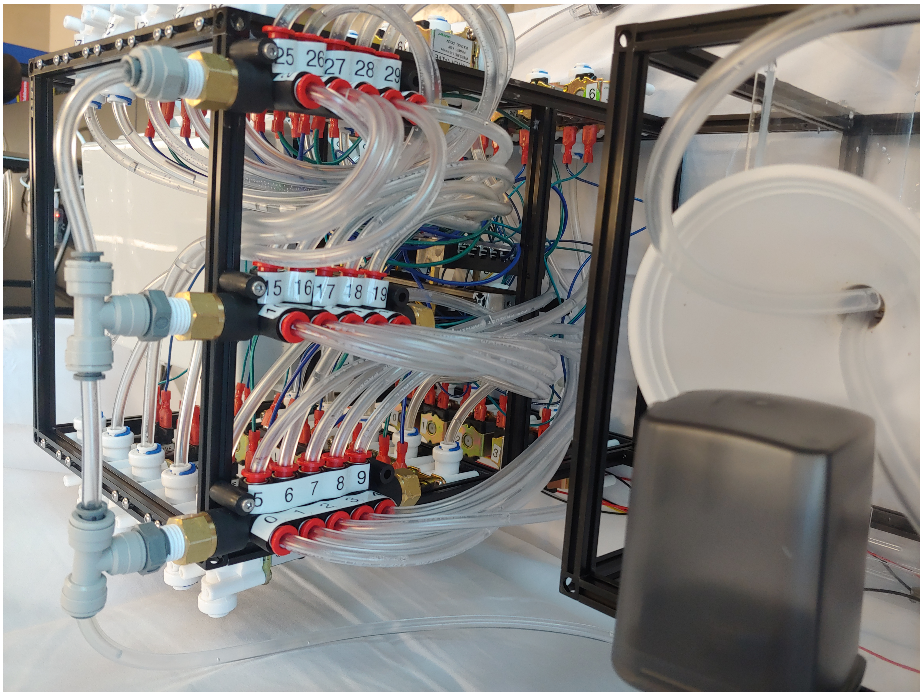


Example of manifolds tubing. ### Become a soldering master

The electronics are probably the most technical part of ZAF+ construction. There is nothing overly complex, but it does require carefull planning and a lot of diligent soldering to connect the different electrical components. We therefore advice that you plan ahead. The electronics are comprised of four different components:

- A Raspberry Pi 3 B+ to run the software and control the electronics.
- Two Arduino Megas (Arduino 2560 R3) microcontrollers for the digital devices.
- Several motor controllers to control the various pumps.
- 16 Relay Module interface board to drive current and control the valves.

To summarize the wiring: the Raspberry Pi is connected directly to one Arduino Mega using a USB cable. The two Arduinos are daisy-chained via a serial connection (the whole design can be extended by daisy-chaining more arduinos). The Arduinos are connected the motor controllers through digital pins (PWM). The Arduinos control the opening of each individual valve through the 16 relay module. A 12V power supply (at least 10A best 20A) provides power to the electronics, except for the Raspberry Pi and the two Arduino Megas powered by the Raspberry Pi 5V.

The following figure summarises ZAF+ electronic connections:

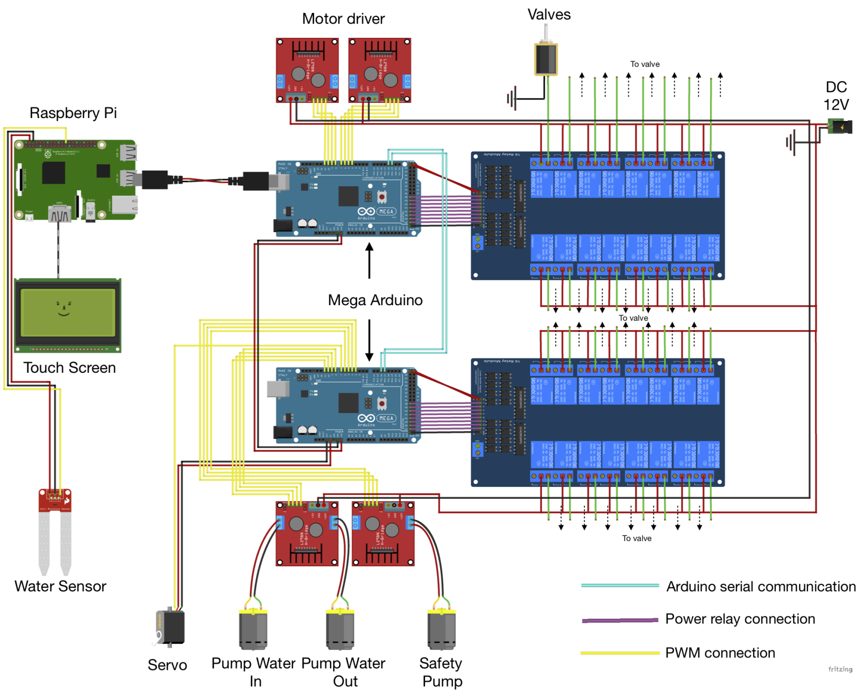


##### Electrical safety box

To avoid any water splashing on to the electronics, which can induce malfunction and/or electrical hazards, all of the core electronic components should be placed inside of a waterproof safety box. We placed all the ZAF+ electronics in the safety box, with the exception of the Raspberry Pi microcomputer and the screen which can be placed far from the water because the only link between that module and the rest of ZAF+ is a USB cable. To ensure safe and easy connections with the external components (i.e valves, pumps etc...), we connected the main electronic components with electrical screw-terminals outside of the safety box. For example, all the downstream connections from the 16 Module Relays are connected to the valves through screw-terminalslocated outside of the box, and which are attached to ZAF+ frame in the valve and tubing compartment. Similarly, the motor drivers are in the box and connected to a screw-terminal, allowing for easy pump connections without opening the electrical safety box.

##### ZAF+ requires many serial connections and proper wiring organisation.

As described in the main electronics figure, there is a lot of wiring in ZAF+, mainly to connect the 16 module relays to the valve through the screw terminal. Therefore, good wiring hygiene is essential. We recommend that you use a solder breadboard to connect and organise the connections from the 16 module relays to the valve terminals.

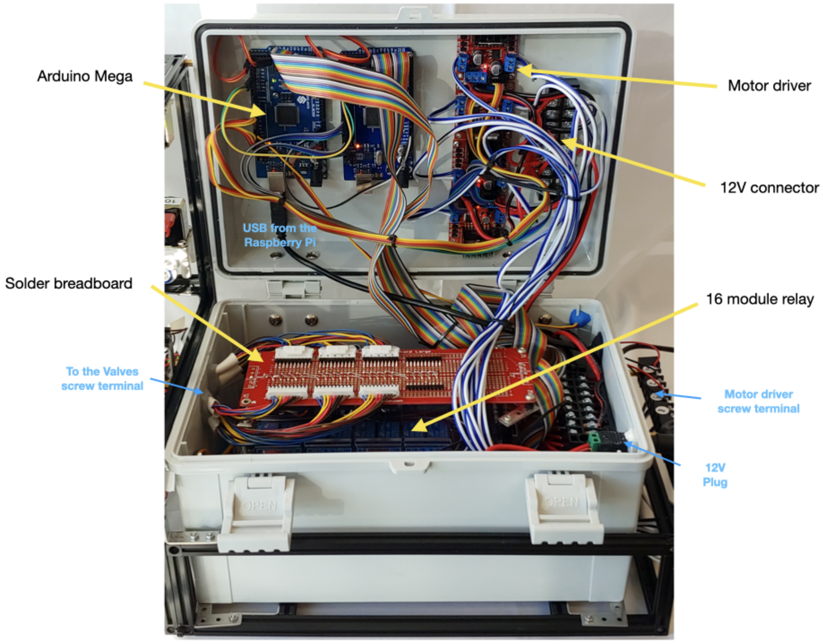


Picture showing the organisation of the electronic box and its different components.

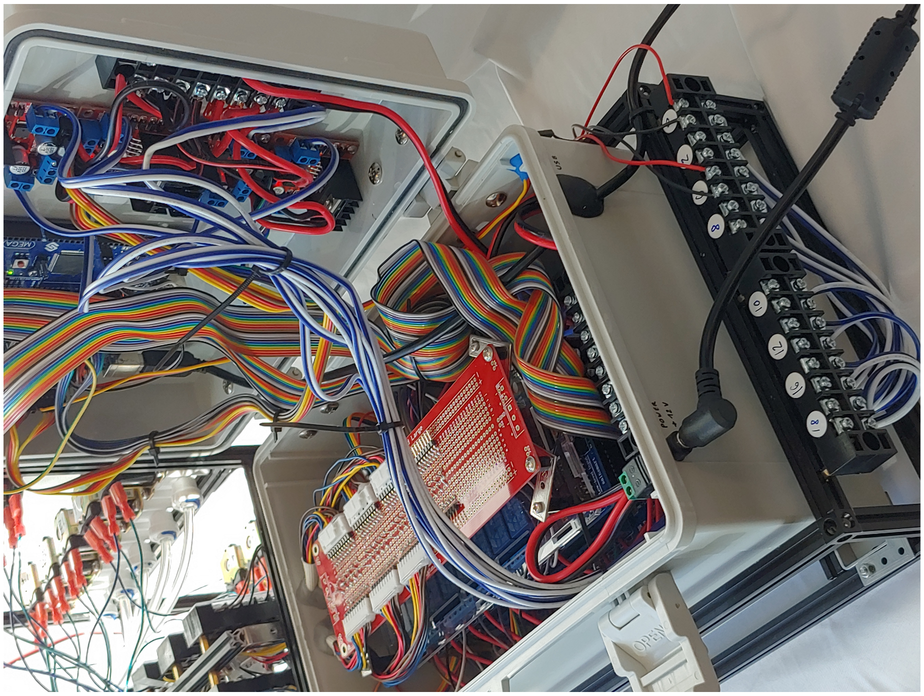

Lateral view of the electronics box.

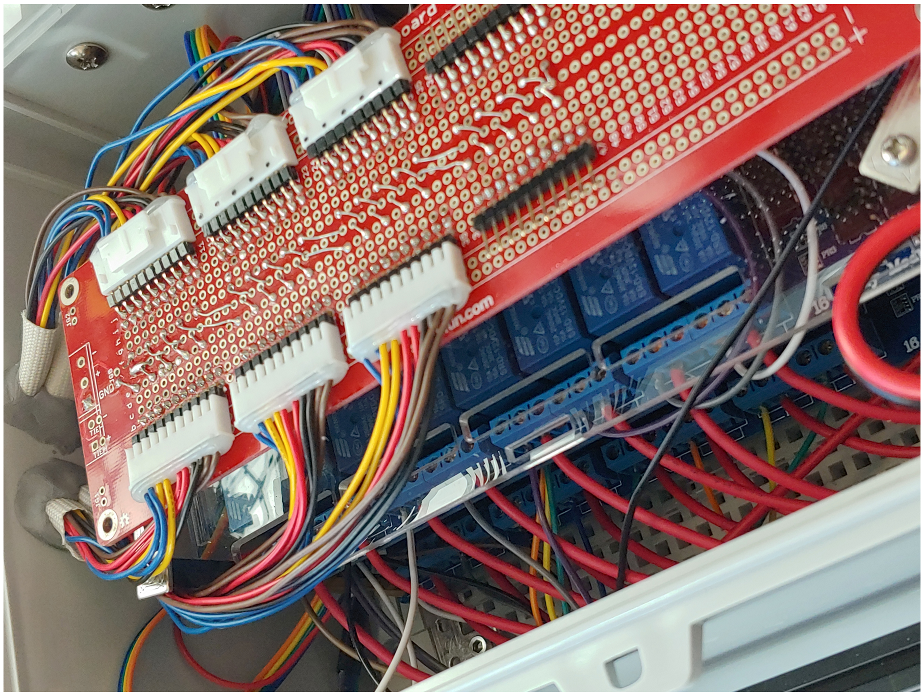

Zoom in view on the 16 module relays and the solder breadboard.

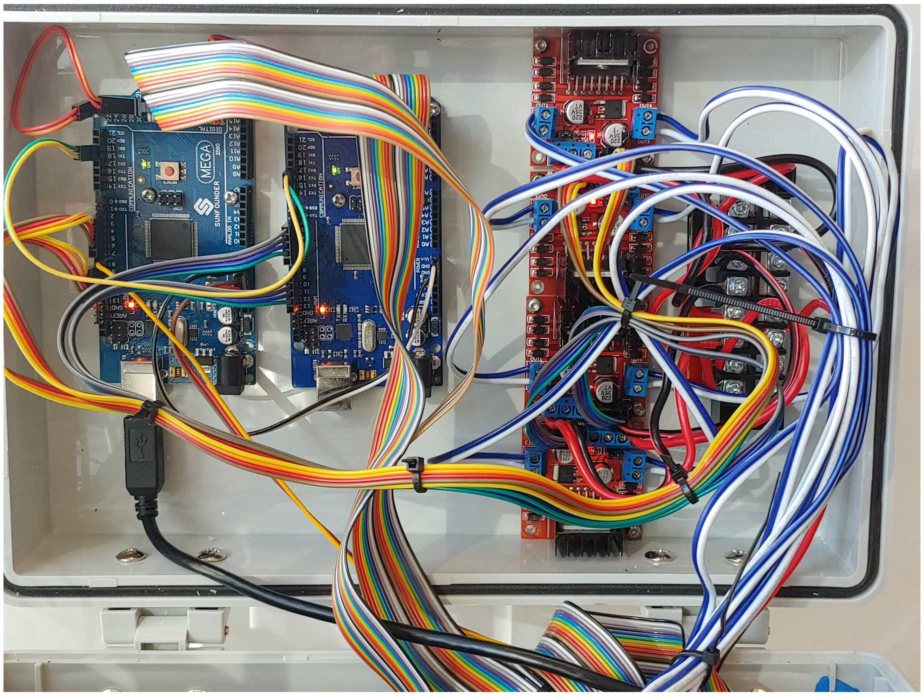

View of the top panel of the box.

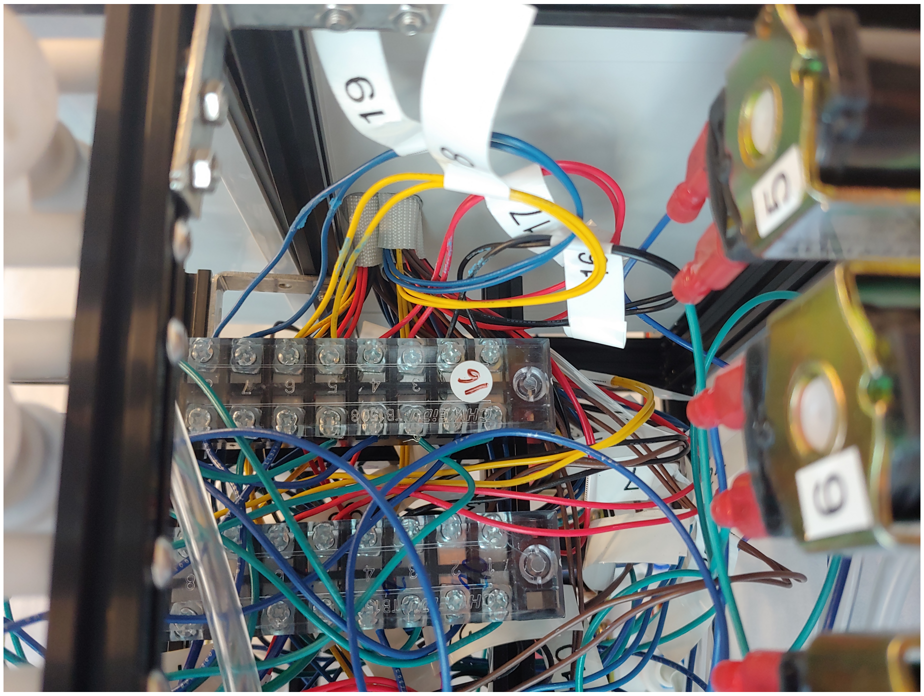

View of the valves screw terminal.

##### Arduino Mega the perfect micro-controller

The Mega Arduino is a micro-controller designed for projects that require many input and output lines. It very easy to use: simply connect it to the Raspberry Pi through a USB cable. You will also need to upload our firmware to your Arduino (see the software section of this wiki). You can daisy-chain two or more Arduino Mega micro-controllers in series to control an almost unlimited number of motors, relays and servos.

You can find the connections detailed in the following diagram.

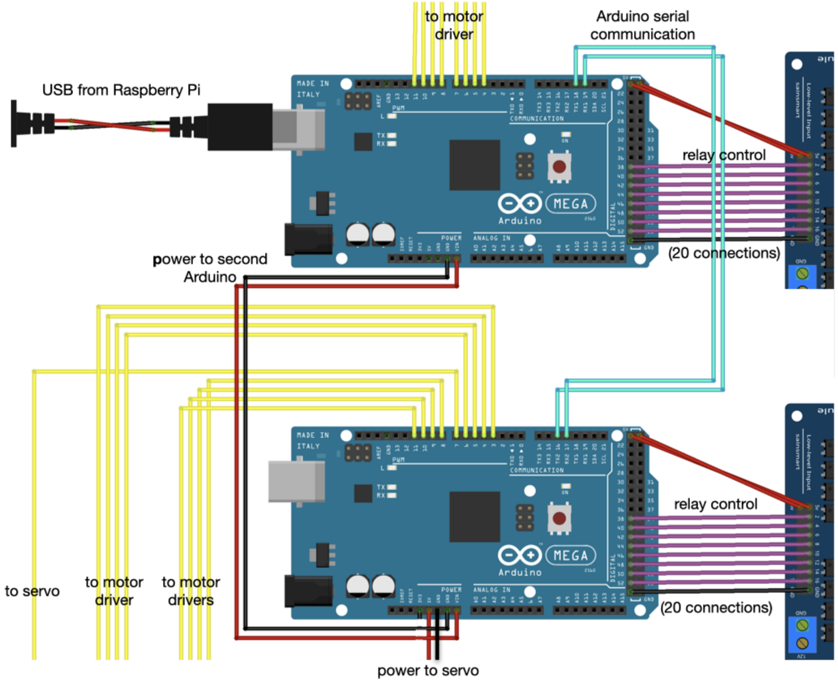

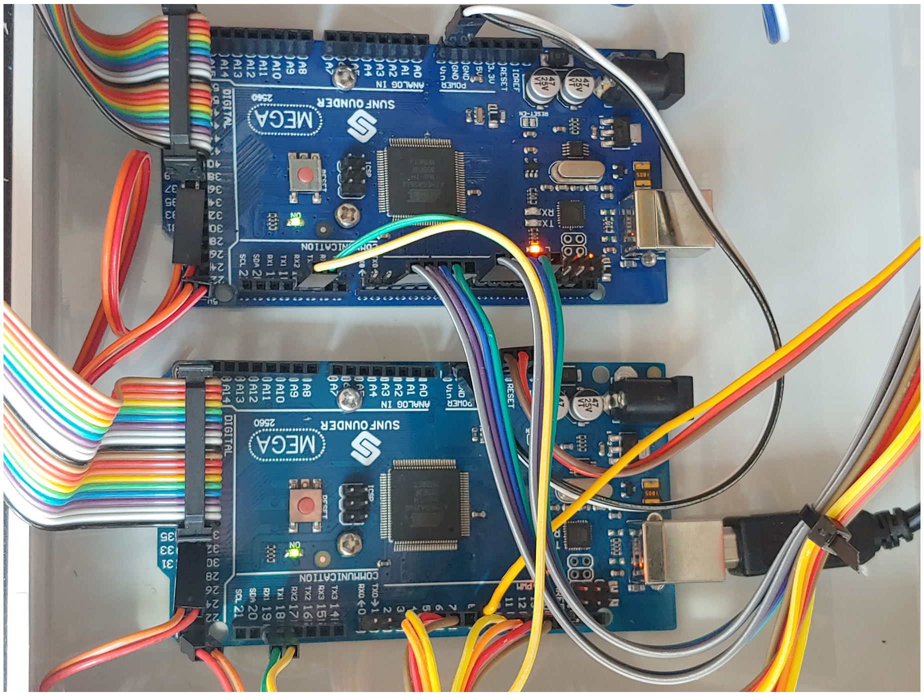


The two Arduino Mega inside the electronic box ### Motor drivers

For a detailed explanation of the connections, please refer to the original ZAF page.

The main difference between ZAF+ and ZAF is that for ZAF+ the PMW connections are coming from the Arduino Mega. In our ZAF+ design we have 4 motor drivers to connect the increased number of pumps (compared to ZAF). This can be easily scaled up if you need a device capable of controlling food distribution for more than 30 tanks.

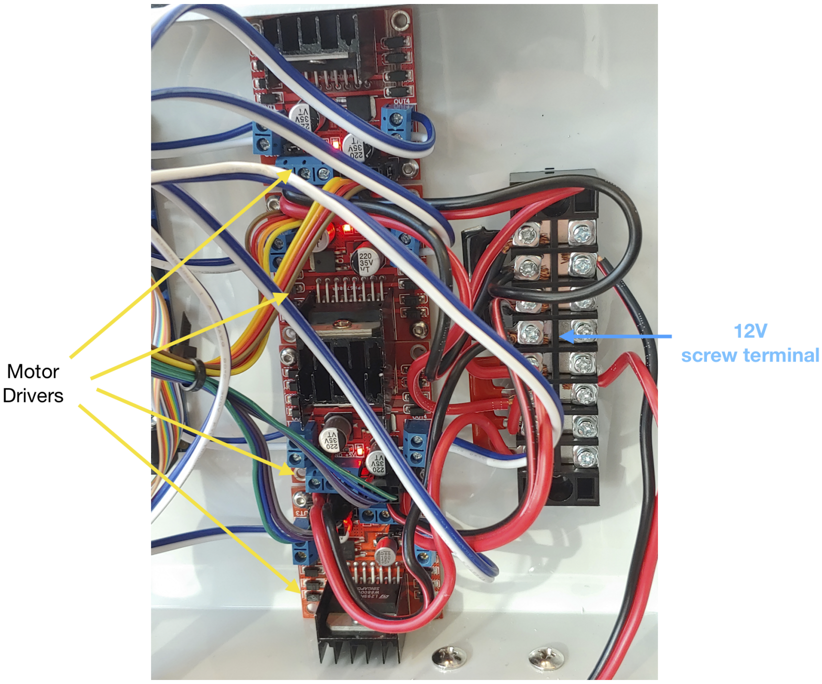


**The 16 relay module to control the valve openings.**

The Relays are control 12V lines that are controlled by 5V digital pins from the Arduino Mega. Each relay board controls 15 12V lines that are used to open 15 corresponding valves. We have 30 tanks in our ZAF+ setup, therefore we need two 16 relay modules. Of course, you can scale up your ZAF+ by adding more relays by daisy chaining more arduinos. The main issue with the relays is the large number of connections. As described herein it is just a matter of planning and good wiring and soldering practices. The 16 relay module inputs and outputs are organised with a solder breadboard connected to each valve outside of the electronic box through a screw terminal.

In the following picture and schema, you can see how to connect and organise the wiring of the 16 module relays.

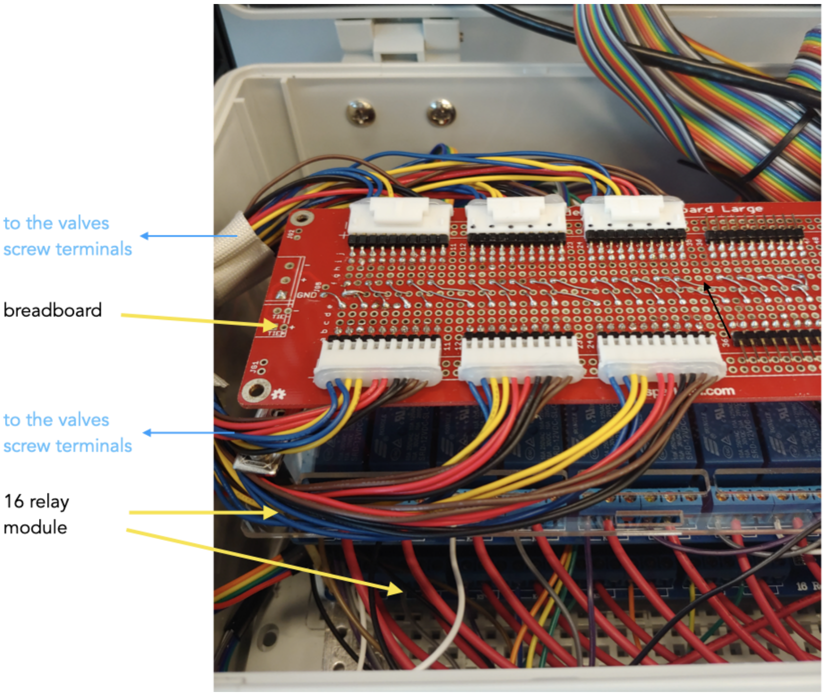
to the valves
screw terminals
breadboard
to the valves
screw terminals
16 relay
module

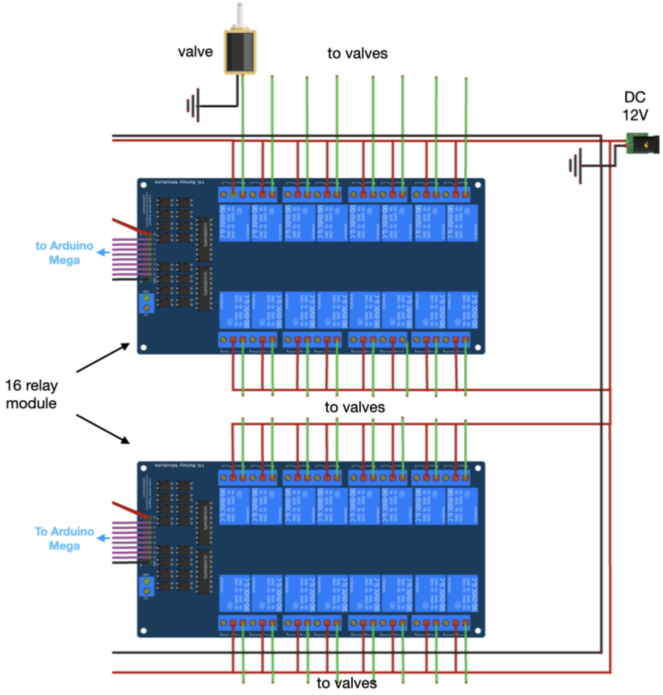
valve
to valves
DC
12V
to Arduino
Mega
16 relay
module
to valves
To Arduino
Mega
to valves

**The computer and screen for the user interface.**

We built a module for the Raspberry Pi 3 B+ and the touch screen using Makerbeam. To set up your Raspberry Pi for easy control, we added a wireless keyboard with a touchpad to the system. Follow this instruction to set up your Raspberry Pi. The touch screen and the Raspberry are connected together with an HDMI cable and a USB/micro-USB cable. For details on the connections and how to fasten them to the Makerbeam structure, please refer to the following pictures.

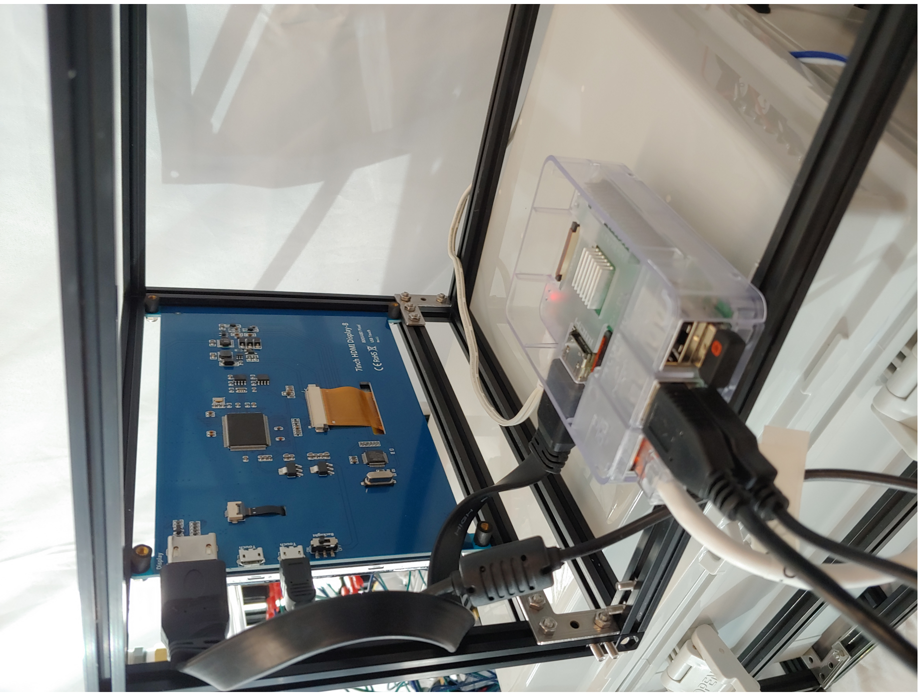

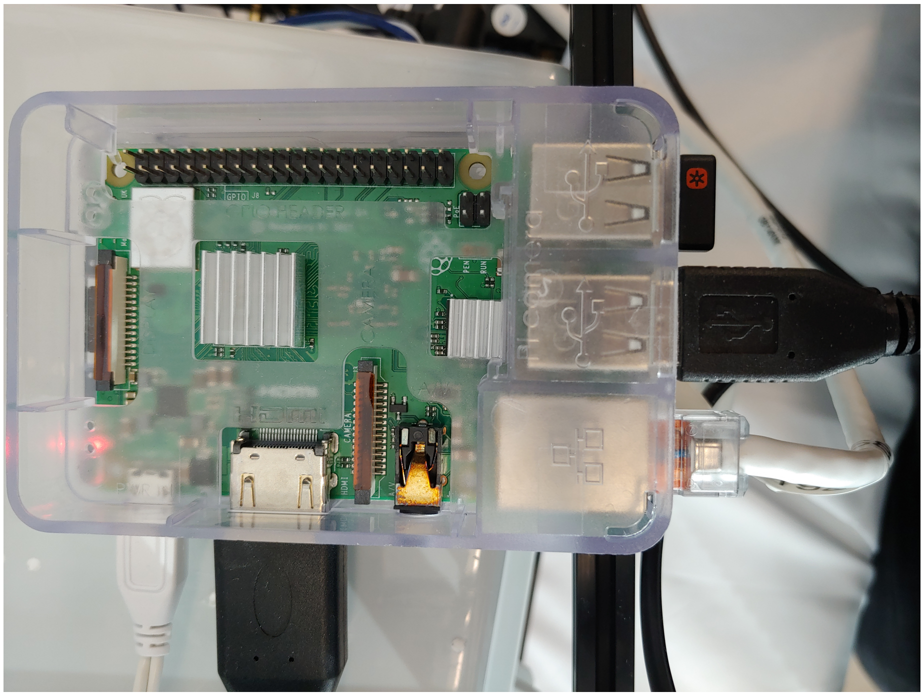

#### ZAF+ Software

The ZAF+ hardware is controlled by an Arduino and a raspberry pi. Arduino is controlling the valves, pumps and food servo. Raspberry Pi communicates with Arduino over serial communication and orchestrate the entire ZAF+ hardware. Drawing below summarize hardware connections and further information can be found in here.

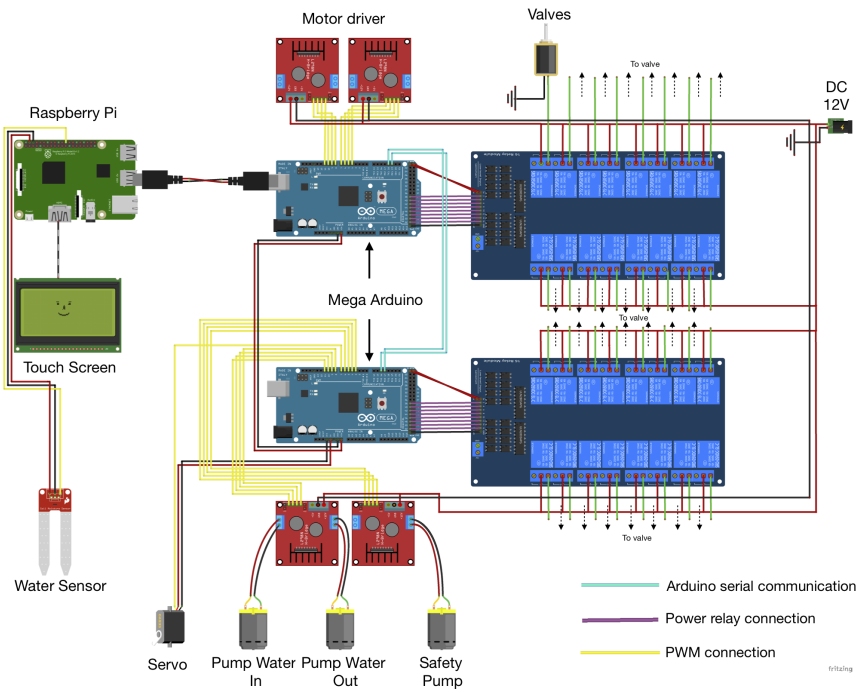


We have developed software pieces for both the Raspberry Pi and the mega Arduinos we used. One can find below the instructions to install and use our software.

- Installation
- How to use

#### How to install?

**Installation on Arduino** The main sketch (controlbox) must be uploaded to both arduino megas. Nothing else needs to be done. If you need help with loading the sketch to your Arduino here you can find a nice tutorial.

**Installation on Raspberry Pi** ZAF+ code is written and tested on only Raspbian Buster >=10.

- First, we recommend you first have a look at the official documentation.
- To get started first download the Raspbian OS and burn it in your microSD card. We recommend doing this with the help of Raspberry Pi Imager which makes this process painless.
- Then you need to get command line access to your Raspberry Pi. You can connect it to a local network via ethernet or wifi. Once connected to the network, you can access a remote terminal using the ssh protocol. A

simpler approach consists in connecting a monitor and keyboard to your Raspberry Pi and opening the terminal.

- After having access to Raspberry Pi terminal, follow the commands listed below one-by-one:

```
# Install dependencies
sudo apt install python3-pyqt5
python3 -m pip install python-crontab==2.5.1 arbol==2020.11.6

# Create the required folder structures
mkdir -p ~/Dev/prod/zaf_data
cd Dev/prod/

# Get ZAF+ software
git clone https://github.com/royerlab/zaf.git
cd zaf
```

- Congratulations! You successfully installed ZAF+ software. ## How to use?

##### Start GUI

After changing the ZAF clone directory in your terminal, you can start GUI with:

```
python3 -m python.gui.gui
```

##### GUI

Using ZAF+ is very simple. There are only three types of tabs in this program: **Dashboard**, **Log** and **Program**. Dashboard displays the overview of the entire program.

On the left half, **Control Panel**, there are two buttons, **Add program** and **Exit ZAF**. You can add a new feeding/cleaning program by pressing **Add program**. To exit ZAF+, press **Exit ZAF**.

On the right half, **Programs**, you can instantly activate or deactivate each program by checking or unchecking the boxes.

The **Log** tab displays all program logs. You can find the record of how and when your programs were executed. You can also find if any error has occurred.

The **Program** tab is where you set all the detailed conditions for feeding and washing.

On the upper left, there are four execute buttons. **Run**: Run the program. **Reset**: Reset the program to the default state. **Duplicate**: Duplicate the current program. **Delete**: Delete the current program.

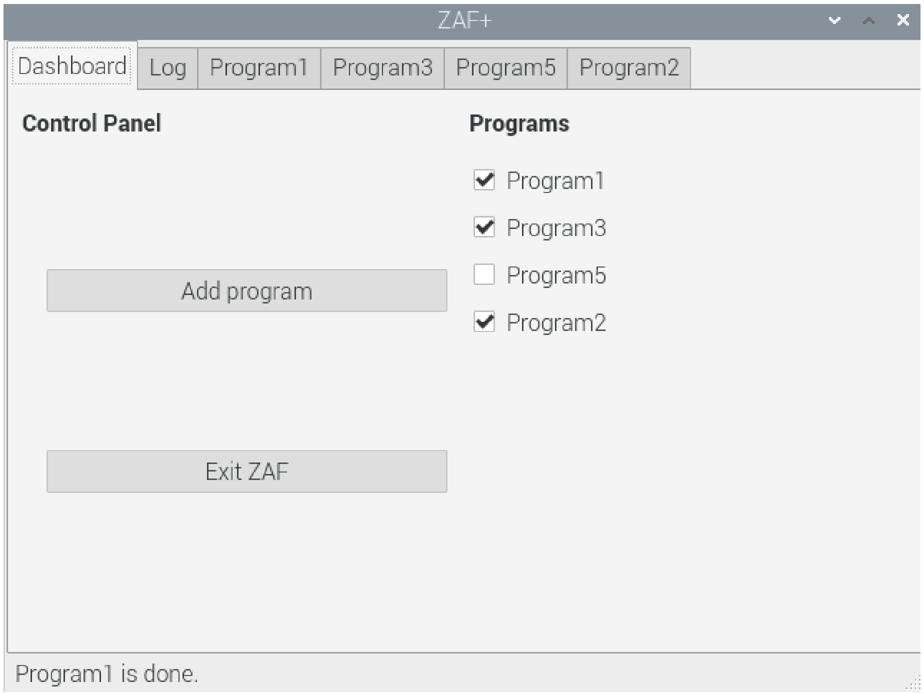


Figure 3: zaf_dashboard_ss

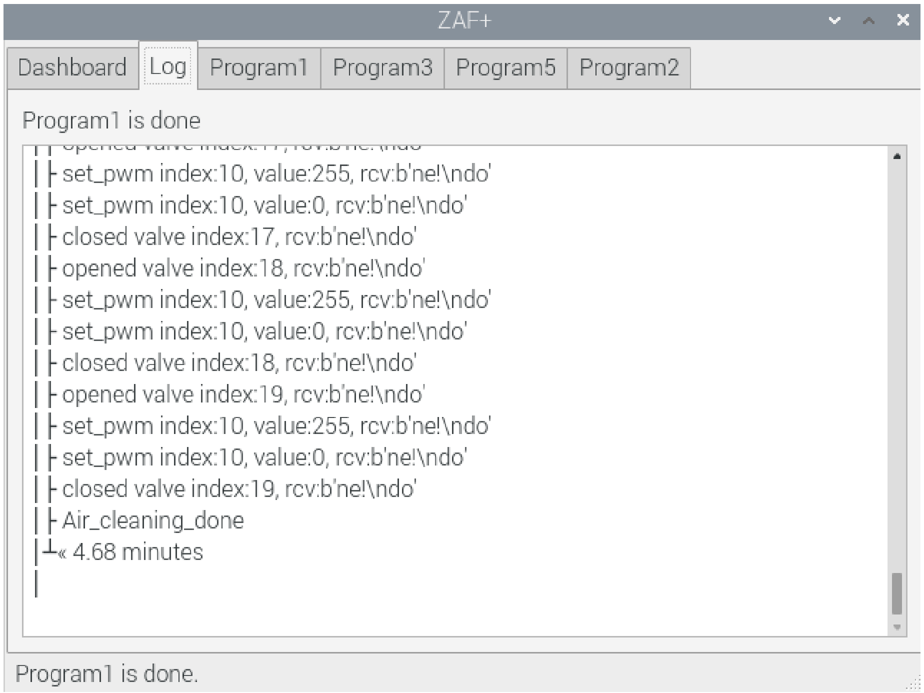


Figure 4: zaf_log_ss

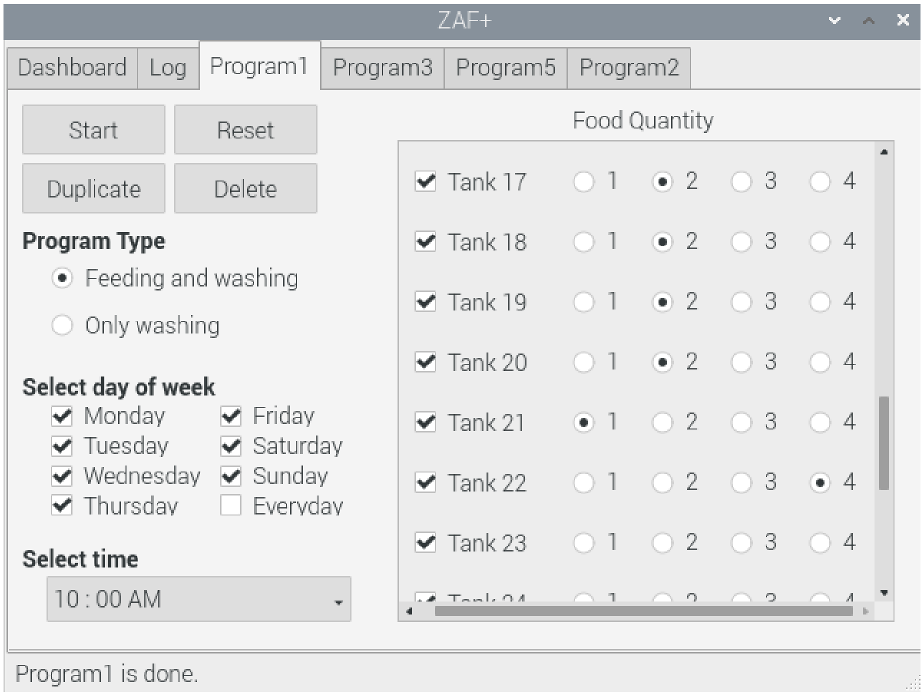


Figure 5: zaf_program_ss

In **Program Type**, you can select two types of program. * Feeding and washing: Feeding followed by washing * Only washing: Only washing but no feeding

In **Select day of week**, you can select the days of week to run the current program.

In **Select time**, you can set the time to run the current program.

In **Food Quantity** on the right side, you can set which tank to feed/wash by checking the boxes and select the amount of food to feed by selecting one of the number from 1 to 4. The higher the number the more food.

###### Persistent Feeding and Washing Programs

ZAF+ GUI saves a small JSON file for each created program and sets the corresponding crontab task to be able to run all the programs no matter GUI is running or not at the scheduled time.

Saved JSON files can be found in the `prod/zaf_data` folder.

#### How to scale up ZAF+?

Here again we can augment ZAF capacity based on the design modularity. A simple possibility is to split the tubes downstream the valves to two or more outputs.

There is also the possibility of mount extra Arduino Mega in the system by connecting them in serial communication and then add a 16 module relay to connect more valves.

To scale up ZAF+, you will need to adapt the code too.

- If you like to add a new pump, you need to first register the new pump in `Context` class we have in `python.zaf_plus.context` submodule.
- Then, you can go to `python.zaf_plus.fishfeed` submodule and use the new pump at any point of the code you like.‘
- For instance if you added a new `water_in` pump, you can adapt your code to use two `water_in` pumps as shown below:

```
Ctx.pwm.setPWM(Ctx.water_in, 0, 4095)
Ctx.pwm.setPWM(Ctx.water_in2, 0, 4095)
sleep(0.5)
Ctx.pwm.setPWM(Ctx.water_in, 0, 0)
Ctx.pwm.setPWM(Ctx.water_in2, 0, 0)
```

If you have have further questions about scaling up ZAF+ feel free to contact us by opening an issue on our repository. ## Necessary Equipments

The following table is indicating the equipments we used to build our ZAF+. Most of the components used are generic and can be replaced by components with similar specs. We added hyperlink on the GitHub wiki to help the user buying/finding the appropriate materials. The only devices that cannot be easily exchanged are the arduino and Raspberry PI, but that is not an issue as these are easily sourced components.

| Components | Parts name | Supplier | Article Number | Number | Unite Price (USD) | Total Price (USD) |
|---|---|---|---|---|---|---|
| Frame | Makerbeam Starter Kit | Makerbeam | 103318 | 1 | 112.25 | 112.25 |
| | 200mm black anodised Makerbeam | Makerbeam | 10090 | 2 | 12.5 | 25 |
| | 300mm black anodised Makerbeam | Makerbeam | 100102 | 2 | 9.75 | 19.5 |
| | Makerbeam corner cube | Makerbeam | 100988 | 1 | 16.95 | 16.95 |
| Servo & Food container | Eheim Automatic feeding unit | Eheim | NA | 4 | 23.47 | 93.88 |
| | Digital Servo | N/A | DS318 | 1 | 16.66 | 16.66 |
| | Magnets | Dymag | 3MM-mix | 1 | 14.99 | 14.99 |
| Pumps & Valve | 12V DC Pumps | Bayite | XX328 | 2 | 29.99 | 59.98 |
| | Solenoid valve quick connect ¼" | Digiten | N/A | 30 | 7.49 | 224.7 |
| | Check valve ¼" | Blulu | 40141600 | 1 | 7.99 | 7.99 |
| Food Mixing | Tube Holder | Asayu | N/A | 1 | 5.99 | 5.99 |
| | Funnel | Karzone | N/A | 1 | 6.99 | 6.99 |
| Safety | Water sensor | DAOKI | TS-VS-292-CA | 1 | 20 | 20 |
| | Minipump | Walfront | 12V DC 6W | 1 | 15.49 | 15.49 |
| Tubing | Soft Plastic tubing ⅜" 50ft | McMaster | 5233k65 | 1 | 49 | 49 |
| | Soft Plastic tubing ¼" 100ft | McMaster | 5233k52 | 3 | 18 | 54 |
| | Push to connect reducer | McMaster | 5779k355 | 1 | 7.30 | 7.30 |
| | manifolds | McMaster | 52045k216 | 3 | 26.64 | 79.92 |
| | T shape connectors | McMaster | 51055k27 | 2 | 4.41 | 8.82 |
| | T shape connectors | McMaster | 51055k195 | 1 | 4.41 | 4.41 |
| Electronics | Waterproof box | Ogrmar | N/A | 1 | 34.99 | 34.99 |
| | Screw terminal X8 | Milapeak | N/A | 1 | 12.49 | 12.49 |
| | Motor Drivers | Qunqi | L298N | 4 | 8.69 | 34.76 |
| | Canakit Raspberry Pi 3B+ | Canakit | N/A | 1 | 69.99 | 69.99 |
| | Solder Breadboard | Sparkfun | PRT-12699 | 1 | 12.49 | 12.49 |
| | LCD Touch Screen 1024X600 | Longruner | LSC7B | 1 | 56.99 | 59.99 |
| | Arduino Mega 2560R3 | Sparkfun | Dev-11061 | 2 | 39.65 | 79.3 |
| | 16X module relay | Sainsmart | 170C20 | 2 | 20.99 | 41.98 |

**Total Price** **1189.81$**

#### Troubleshooting

- In the following table we present solutions to the most common minor problems we encountered during ZAFs construction and during their operations.
- Overall ZAF has been tested for a period of 9 months and ZAF+ during 10 months. We never experiences any major problems!
- We invite all ZAFs developers/users to report problem, bugs or comments to use ZAF Github issues.

| Problem | Cause | Solution |
|---|---|---|
| - Burning of electronic components (ZAF specific). | - Voltage ripples due. | - Add 1000uF capacitors in parallel to the motor terminals |
| - Food distribution is non homogeneous (ZAF specific). | - tubing issues. | - All the tubes should be same length<br>- One tube might be pinched or bent. |
| - All the pumps connected to a motor controller (LN298) are not working. | - Wiring problem. | - Check the T splice connector wiring connections |
| - A lot of food remains in the food preparation tank. | - bad pumping. | - Check the tube position inside the food preparation container, it must be at the bottom.<br>- Possible pump malfunction, change the pump. |
| - No food going inside the food preparation tank. | - Food delivery is clogged. | - Clean the funnel and the food dispenser output. |
| - Water leak at the pump output. | - Tubing connections with the pump. | - Change the zip tie and eventually cut the tip of the damaged tube. |
| - Dirty fish water feeding . | - Overfeeding. | - Reduce the food quantity delivered by i. Adjusting the software, ii. Adjust the food dispenser closure. |
| - Tubes are getting dirty quickly. | - Cleaning is not well done. | - Add more cleaning programs. |
| - The mixed water/food is not going out of the tube | - The pump is not running correctly and or tube are too long | - Possible pump malfunction, change the pump.<br>- Increase the pump running period in the code source. |
